# Genebank genomics allows greatly improved taxonomic correction for *Capsicum spp*. accessions using a novel automated classification method

**DOI:** 10.1101/2022.11.09.515845

**Authors:** M. Timothy Rabanus-Wallace, Nils Stein

**Affiliations:** University of Melbourne, Victoria, Australia; Leibniz Institute of Plant Genetics and Crop Plant Research (IPK), Seeland, Germany; Center of integrated Breeding Research (CiBreed), Georg-August-University, Göttingen, Germany

**Keywords:** Capsicum, Pepper, Genebank, Genomics, Taxonomy, Classification, Species

## Abstract

To maximise the benefit of exploiting genebank resources, accurate and complete taxonomic assignments are imperative. The rise of genebank genomics allows genetic methods to be used for this task, but these need to be largely automated since the number of samples dealt with is too great for efficient manual recategorisation, but no clearly optimal method has yet arisen. A recent landmark genebank genomic study sequenced over 10,000 accessions of peppers (*Capsicum spp*.), for which the exploitation of genebank material is of huge commercial, cultural, and scientific importance. This study resulted in precisely the type of dataset that will, in coming decades, be likely be produced for hundreds of plant taxa. The long-appreciated difficulties of pepper taxonomy are evident from the many obvious misclassifications noted in this and other studies, providing a perfect opportunity to simultaneously advance methods development in the area, to correct many genebank taxonomic assignments of pepper accessions, and to provide insights into pepper taxonomy in general. This paper aims to achieve these goals using an approach that combines several ideas from standard classification algorithms to create a highly flexible and customisable classifier that performs favourably when compared with key alternative methods. The various characteristics of different methods are discussed, and possible sensible alterations to pepper taxonomy based on the results are proposed for discussion by the community.

## 1 Introduction

The importance of genebanks for the collection, maintenance, and exploitation of plant genetic diversity has been recognised for well over a century (Mascher et al., 2019). Accurately characterising and describing this genetic diversity is key to its exploitation. The rise of genebank genomics, and associated herbarium genomics (Bakker et al., 2020), has liberated this effort from its former dependence on measurable phenotype-based approaches such as morphology, phenology, cytogenetics, and chemical assays, allowing direct interrogation of the genetic code of, in notable recent cases, tens of thousands of accessions.

Linnean taxonomic classifiers have always served a useful purpose in this regard, but the Linnean system of nested classifiers with defined levels does not map neatly onto the realities of evolution and reproductive biology, and hence the classifications do not always reliably reflect true biological divisions (such as reproductive boundaries or cladistic monophyly). This generates much discussion, especially in cases such as hybridising species, ring species, or instances of horizontal gene transfer, where the Linnean system struggles most to act as a meaningful classifier. Plants, being particularly prone to such cases, often suffer taxonomic ambiguity (Rouhan et al., 2018). It is generally agreed, nevertheless, that taxonomic classifiers have great utility, functioning at a very minimum as shorthand labels for groups of genetically- or morphologically-similar individuals, whether or not they are truly monophyletic or reproductively-isolated groups. Typically, researchers working with members of a taxon gain an understanding of how well the assignments reflect, or fail to reflect, biological factors of importance to their research. Ultimately, taxonomic assignments may variously function as reliable and convenient indicators of reproductive boundaries, crossability, phenotypic and physiological traits, cytology, agricultural value, and geographic origin, just to name a few.

Genebank genomics—the sequencing of genebank accessions *en masse* for genomics applications such as population genetics, association studies, and enhanced genebank curation—is a rapidly growing field, with recent landmark studies sequencing several thousands or tens of thousands of accessions. Accurate and complete taxonomic assignments are imperative not just for maximising the benefit of genebank resources, but also to ensuring their exploitation, since researchers of all kinds have a tendency to focus on germplasms with complete passport information (Meyer, 2015), resulting in neglecting a large fraction of the stored materials where documentation is less complete but might anyway be very favourable for breeding.

One function of these studies is to reveal and attempt to correct taxonomic assignments (or to add assignments where they are missing). The problem of automating taxonomic assignment is not peculiar to genebank genomics. For example, metagenomic studies face the problem of estimating taxon abundances using a mixed sample (Thomas et al., 2012), while bacterial sequence databases ensure taxonomic assignment consistency by automatically checking new submissions against a reference database of type assemblies (Ciufo et al., 2018). Genebank taxonomic correction is, however, a fairly novel task, consisting of using genetic data from a large number of accessions that already have assignments recorded in their passport information, then using those assignments to ascertain how these assignments generally map onto the genetic diversity space of the accessions, then identifying accessions that do not conform with this mapping. The data one corrects is also the source of information by which correction is achieved. This gives the procedure something in common with both smoothing and clustering algorithms. The process has occasionally been done manually where very few corrections have been called for (e.g., Singh et al., 2019; Milner et al., 2019). Several automated methods (summarised aptly by van Bemmelen van der Plaat et al., 2021) for taxonomic assignment, several borrowed from literature on species barcoding and metabarcoding (e.g. using the *cox1* gene sequence), have been suggested and tried in comparative studies (van Bemmelen van der Plaat et al., 2021; Austerlitz et al., 2009; Weitschek et al., 2014), with the finding ultimately being that there may be no best all-round tool, though k-nearest-neighbours (in particular with *k* = 3) classifiers are perhaps the best candidate and generally performed well on test datasets despite their sensitivity to parameter selection and to variation in sample sizes between different taxa in the sample.

A recent study on *Capsicum* accessions from a pan-Eurasian selection of genebanks (Tripodi et al., 2021) featured over 10,000 accessions in total, valorising the dataset with investigations into genetic predictors of agriculturally-relevant traits, and the prominent influence of human trade and migration on the distribution of peppers around the world. Domesticated peppers have supreme economic importance, their visual and gustatory diversity, including the characteristic pungency of many varieties, making them a staple component of cuisines from across the globe. Close to 40 million tonnes of peppers were grown in 2020, with Africa, Eurasia, and the Americas all contributing significantly to the total global yield (FAOSTAT, accessed 11/8/2022). These cultivated peppers fall mainly within the species *C. annuum* (e.g. bell pepper, cayenne pepper, jalapeno), with significant contributions from *C. frutescens, C. chinense*, and *C. pubsescens*, and minor contributions from a host of other semi-domesticated and wild divisions such as *C. baccatum, C. chacoense*, and *C. eximium*. The difficulties involved in *Capsicum* taxonomy are reviewed in detail by Esbaugh (2019), who emphasises that inconsistent assignments frequently occur, with boundaries between even major established *Capsicum* divisions such as *C. frutescens* and *C. chinense* often being unclear. This, notes Esbaugh, is partially a consequence of a lack of firm criteria on which the divisions are based, though fruit shape, pungency, chromosome number, and more subtle phenotypic characters such as the morphology of the corolla, have all played important parts in motivating taxonomic delineations. With large and active communities of breeders and researchers accessing genebank pepper accessions and working with them to produce new pepper varieties with desirable agricultural properties and/or cosmetic/culinary appeal, updating the taxonomic assignments of accessions to reflect true genetic demes is a highly desirable and productive pursuit.

The Tripodi et al. (2021) study applied a clustering-based method to suggest alternative taxonomic classifications for putatively-misclassified *Capsicum* accessions: briefly, accessions were clustered using a UPGMA algorithm with an arbitrarily-defined distance cutoff, and minority members of clusters dominated by a single majority taxonomic assignment (80% of members, in this case), were all given the majority assignment. This method suffers from several flaws, most critically, sensitivity to over-representation of members of one taxon, which is also suffered by the methods reviewed by van Bemmelen van der Plaat et al. (2021): Taxa comprising many samples are by default more likely to dominate clusters, no matter what the cluster size. The capsicum dataset, being dominated by the species *Capsicum annuum*, is particularly vulnerable to methodological biases of this kind. In addition, the composition of clusters is somewhat arbitrary, and results would be expected to differ if the sampled taxa were altered. Furthermore, appropriate visual inspection methods reveal that many taxa clearly associated with a distinct deme having some obvious majority assignment were not given the obvious assignment, most likely owing to their falling—mainly by chance—into a heterogeneous cluster. While better optimisation of the cluster size cutoff and majority cutoff parameters may have helped to alleviate this effect, it is ultimately a result of the clustering-based method itself, and without employing any visualisation tools for the specific purpose of assessing how intuitive the assignment results were, the study ultimately left much room for improvement on the taxonomic assignment front (see section on visualisation in methods; fig. 1).

**Figure 1.**
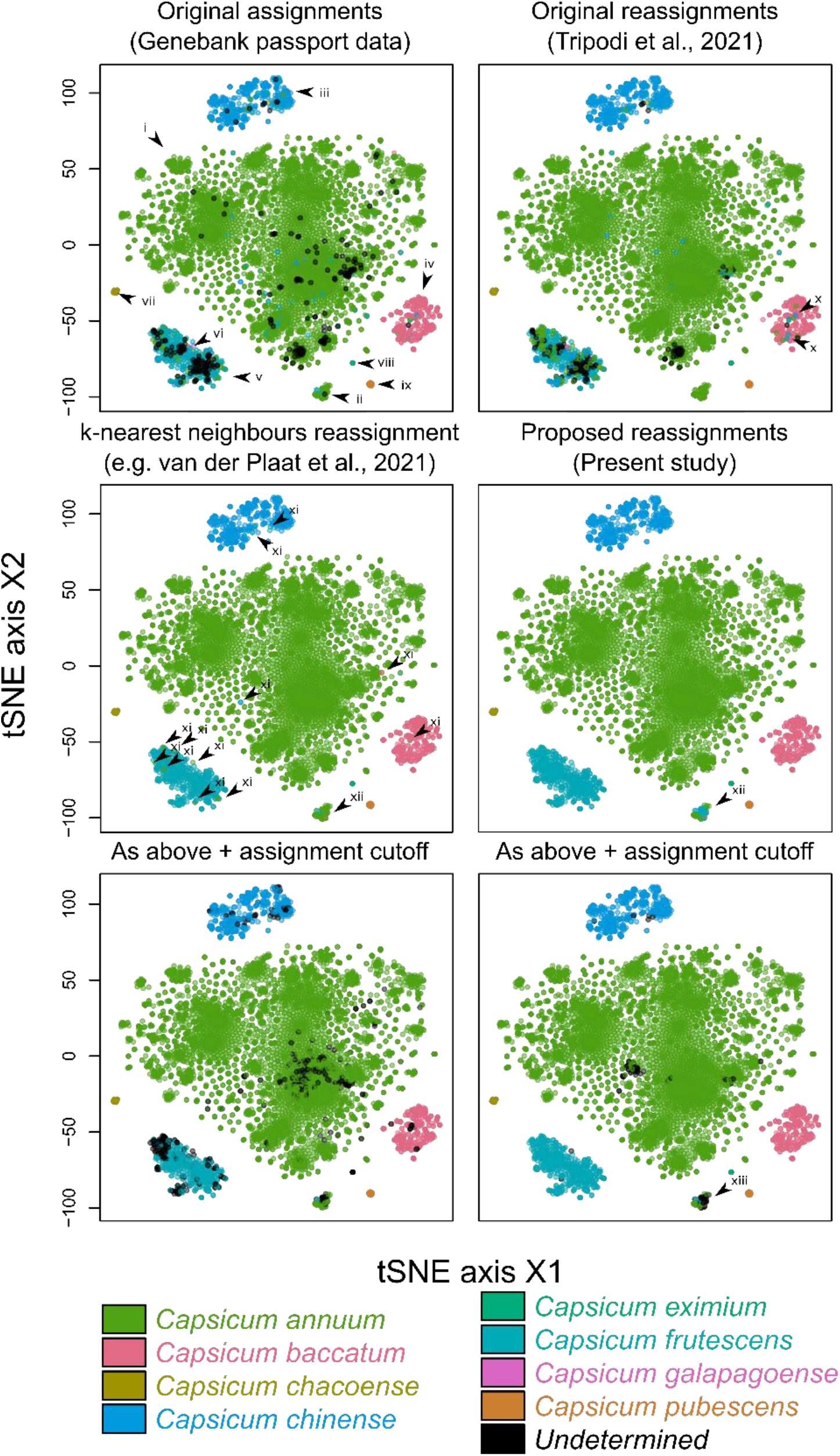
t-SNE plots (calculated from IBS distances, perplexity = 30) representing the relationships among the G2P-Sol pepper accessions used in the study (trimmed dataset—see Methods) and their classifications under different schemes. Markers labelled with Roman numerals are referred to in the main text. For k-nearest neighbours, *k* = 6, which I judged to be the best result possible (see Supplemetary Materials for a range of values). At *k* = 7, *C. eximum* (marker ix) are reassigned as *C. chacoense*. The unassignment cutoff r is set to 4/6. Parameters for the kernel-based method are *𝜆* = 320, *d* = 0.2, *r* = 0.66, *k* = ∞.

**Figure 2.**
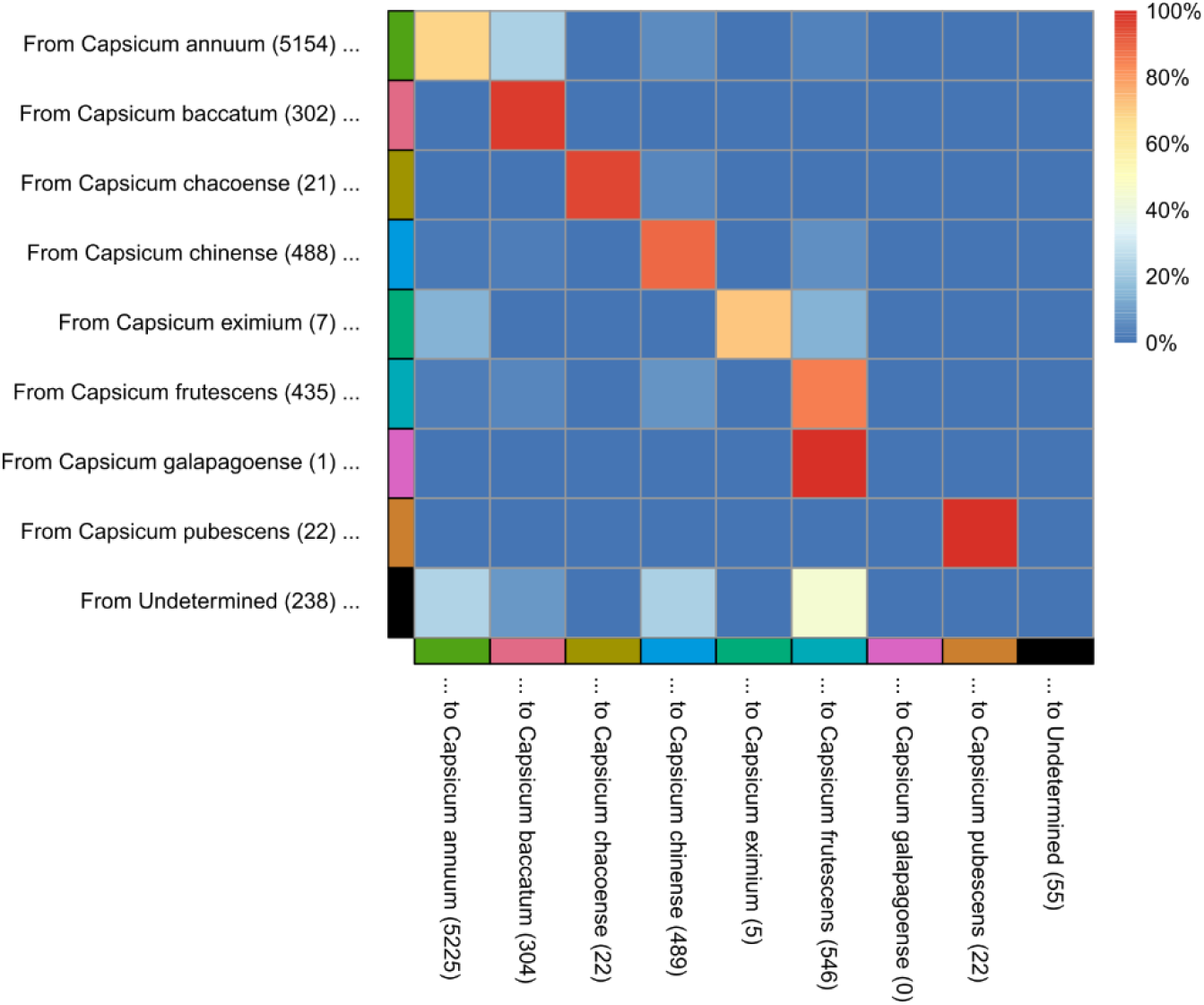
Reclassification rates for *Capsicum* divisions in the dataset using the kernel-based method presented in this study.

We present here a method of automatically producing taxonomic assignments for genebank *Capsicum* accessions that aims to align with human intuitions about their intended use, and potentially to possess broader utility for genebank genomics projects.

## 2 Methods

### 2.1 A new, hybrid approach

The method employed is distinct from its alternatives in its motivation to provide taxonomic assignments that make a compromise between the multiple ways that members of a taxon are expected to be similar: morphologically, cytologically, genetically, phenologically, reproductively, evolutionarily, and so on. To do so, We recognise that taxonomic assignments are given based on various measures of similarity that either suit the conventions of the research community, or which are simply convenient. These, We argue, ought to be respected, but can be occasionally modified to ensure better conformity with the expectation that membership in a taxon entails genetic similarity with other members of the same, but does not exclude the possibility of paraphyletic groupings.

We therefore suggest a method that examines each individual in the dataset and suggests an assignment based on the assignments of the individuals nearest to it, based on identity-by-state genetic distance, similar to the k-nearest neighbours approach that achieved equal-best all-round success in the comparative study of van Bemmelen van der Plaat (2021), by creating a hybrid method that combines k-nearest-neighbours with a variant of the kernel-based Parzen-Rosenblatt Window approach: The suggested assignment of each individual in the dataset to a taxon is made based on the assignments of all other individuals within a predefined distance *d* from the focal individual. Neighbours with an “undetermined” assignment are ignored. The assignment of each neighbour contributes weight to a potential assignment of the focal individual. This weight depends on two factors. Firstly, the IBS distance of the neighbour from the focal individual, which decays exponentially, i.e., by a factor of *e*^−*𝜆* ∙ distance^, with some predefined constant *𝜆* controlling the decay rate. Each potential assignment is given a total score by summing the weights of neighbours with that assignment. Secondly, the scores for each potential assignment are (optionally) normalised by the inverse of the frequency of neighbours bearing that assignment (i.e., individuals within d of the focal), a means to alleviate bias towards oversampled taxa. For instance, for a focal individual with ten neighbours within *d*, 4 from taxon A and 6 from taxon B, the total score for A will be scaled by 10/4 and the total score from B scaled by 10/6. Individuals receive no contribution from themselves, meaning their assignment is ultimately entirely dependent on that of their neighbours. Where the highest-scoring potential assignment out of all the possible assignments for an individual proportionally greater than some pre-defined cutoff *r*, then this assignment is suggested for the focal individual, otherwise an “undetermined” assignment is suggested. A parameter *k* for excluding all but the *k* nearest neighbours is also incorporated. Running the method with *𝜆* = 0, *d* = ∞, and the weighting of neighbours inversely to their taxon’s frequency disabled, is equivalent to the k-nearest-neighbours approach.

### 2.2 Visualisation

The challenge of assessing how well the results meet the goal of retaining established taxon groupings while ensuring individuals in a taxon are genetically similar requires a visualisation method optimised to display genetic similarity and dissimilarity among samples in two dimensions, with minimal loss of information. T-SNE, a dimensionality reduction and visualisation method that specifically aims to optimally preserve inter-item distances, was judged ideal for this purpose. Visual inspection of t-SNE plots (Supplementary Data) was used to select appropriate values of *𝜆, d, r*, and *k* (see figure 1).

### 2.3 Comparing methods

For the sake of comparison, the results of the UPGMA-clustering-based method of Tripodi et al. (2021) were also inspected, and an additional reclassification was done using the k-nearest neighbours method recommended by van Bemmelen van der Plaat (2021). Statistics comparing the outcomes of the methods were generated using the deduplicated dataset, omitting the mixed cluster (fig. 1, xii), with parameters for the kernel approach set as per figure 1, including the unassignment cutoff.

### 2.4 Data curation and treatment of duplicates

Before creating assignments, the dataset was trimmed to include only individuals with valid IBS scores (some IBS scores are marked N.A. in the dataset owing to, for instance, insufficient data), and which were not identified as duplicate accessions in the study of Tripodi et al. (2021). In the case of duplicate accessions, one was randomly selected to represent the whole set of duplicates. The final assignment of the chosen accession was then transferred to the rest of the set.

## 3 Results and Discussion

### 3.1 Passport classifications

Visual comparison of the three methods of classification show marked differences in efficacy, with assignments using 3-nearest-neighbours and the present method being far more intuitive and corresponding with demes evident in the t-SNE plots (fig. 1). The passport classifications visualised by t-SNE shows that *C. annuum* refers to one major deme (fig. 1i; henceforth roman numerals refer to markers in figure 1), although it appears to dominate a second smaller deme that contains a mixture of taxa (ii). *C. annuum*’s dominance in this secondary deme may, however, owe only to its being so well represented in the dataset compared with other taxa. Other important demes are dominated respectively by *C. chinense* (iii), *C. baccatum* (iv), and *C. frutescens* (v). Members of this *C. frutescens* deme are frequently classified as *C. annuum*, and also includes the dataset’s only instance of *C. galapagoense* (vi). Occasional unintuitive assignments can be seen in all the major demes but seems especially common in the *C. frutescens* deme, likely due to some ambiguity in morphology-based classification, perhaps explaining the large proportion of undetermined accessions in this deme. Other minor taxa assignments *C. chacoense* (vii), *C. baccatum* (viii), and *C. eximium* (ix) appear in this dataset to represent genetically separated groups well deserving of a distinct designation.

### 3.2 UPGMA-clustering-based reclassifications

The Tripodi et al. (2021) UPGMA-clustering-based reclassifications make only minor improvements. A good proportion of taxon-undetermined accessions are given assignments but many remain, and while some reclassification appears to remove non-intuitive assignments, many of these remain too. However, some of the reassignments appear to actually introduce new errors, for instance within the *C. baccatum* deme, where small clusters that by chance capture closely-related misclassified taxa within a larger deme enable members of the dominant taxon to be reclassified as the minority taxon (x).

### 3.3 K-nearest neighbours

The k-nearest-neighbours approach makes significant improvements both to assigning undetermined accessions and to correcting unintuitive assignments, but is very much subject to a problem analogous to that just described for the UPGMA-clustering-based method above. This problem persists even when the neighbour count is increased, but before it eliminates all remaining undetermined or unintuitively-classified accessions, it begins to reclassify minor taxa with few representatives such as *C. eximium* (ix) as members of clearly-genetically-distinct taxa (in this case, *C. annuum* or *C. pubsescens*, depending on *k;* Supplementary Data). Overcoming this problem by introducing weightings based on distance and neighbour assignment frequency was a major motivation for the kernel-based method introduced in this publication.

### 3.4 Kernel-based approach (present study)

The kernel-based method successfully overcomes these problems, unifying assignments in the one-taxon-dominated demes, including those with very few members, with, in the authors’ judgement, no unintuitive “out of place” assignments (fig. 1). And the number and nature of the reassignments made to achieve this level of congruency is substantial. Compared with the UPGMA-clustering method, the kernel approach suggested approximately three times as many assignment alterations (2.4% vs 7.5% of all accessions). Of all the cases in which the kernel method suggested alterations, only 30.1% of the final assignments matched those suggested by the clustering method. And where the clustering method decreased the number of undetermined assignments by 31.8%, the kernel method gave 98% of undetermined accessions an assignment. The best results were attained with *k* = ∞ (effectively disabling the *k*-nearest-neighbour functionality). The inverse weighting of neighbours by assignment frequency proved essential for achieving this level of accuracy without causing genetically-distinct taxa with low sample numbers to be collapsed into other taxa. Similarly, manipulation of the decay parameter is particularly important to strike a balance between under-sensitivity and overgeneralisation (Supplementary Data). The ability of the kernel-based method to give the user very fine control over a number of continuous parameters (c.f. discontinuous parameters like *k* in a k-nearest-neighbours approach) is responsible for this essential flexibility, which we believe will make the method a valuable contribution to achieving the equivalent tasks across diverse datasets. This includes continuous control over the parameter *r*, describing how much a potential assignment’s score must dominate for the focal individual to be given an assignment. The equivalent to *r* in k-nearest-neighbours is discontinuous since it must designate some integer number of neighbours required to possess a shared assignment. The lower four panels of figure 1 demonstrate how this allows the kernel-based method to better chose a value of *r* that eliminates unintuitive classifications but induces minimal extra undetermined samples that may require manual work to reclassify. Both take the lowest *r* possible to remove obviously unintuitive assignments, but the kernel-based method is able to induce fewer unassigned calls within large demes where the assignment seems obvious.

### 3.5 Further comments on the kernel-based approach and implications of the results for *Capsicum* taxonomy

The reassignments suggested by the approach support the existence of all pepper species featured in the study with the exception of perhaps *C. galapagoense*, though little can be concluded since it is here represented by a single accession. The limited number (n = 7) of *C. eximium* accessions in the study had a markedly high tendency to be putatively-misclassified *C. annuum* and *C. frustescens*, and *C. baccatum* included many putatively-misclassified *C. annuum*.

A key question raised by the observations of this study but no doubt broadly applicable to genebank genomic and taxonomic studies, is the appropriate treatment of genetic clusters that contain mixed species assignments, the best example of which here is marked ii in figure 1. A strength of the kernel-based approach is that, owing to frequency-based normalisation of potential taxon scores, the mixed cluster is not as susceptible to being subsumed into its most common member simply owing to that member being better represented in the dataset, as occurs with the UPGMA-clustering-based method (fig. 1), and k-nearest-neighbours for many larger values of *k* that would otherwise make for sensible choices (Supplementary Data). By alleviating this bias, the mixed cluster retains its mixed character and moreover, when an assignment cutoff is imposed, it is quickly flagged to the investigator that the cluster is taxonomically ambiguous, owing to the prevalence of unassigned samples within it. I defer to the *Capsicum* taxonomy community to discuss the merits of assigning members of this cluster a specific, unified designation.

Equally important as parametric flexibility and ambiguity handling, is the visualisation method used to assess how well chosen parameters achieve sensible results, especially in cases where the actual genetic structure of the population and distribution of taxon assignments over the genetic diversity space is initially unknown. Naturally every possible choice of visualisation tool has tradeoffs and one must be cautious of, for example, the distortions that can be induced by dimensionality reduction.

Such distortion may explain why some accessions are persistently given assignments that surprisingly do not match those of the accessions clustered around them in the plot (Supplementary Data). Tripodi et al. (2021) present some evidence that suggests highly-heterozygous early-generation hybrid individuals may be a key cause of this taxonomic ambiguity, and indeed the challenge of heterozygosis in obligate outcrossing species means that future studies employing methods such as that presented, which require a distance matrix, will need to consider more sophisticated distance measures than IBS before these methods can be applied.

### 3.6 Outlook

Solving the problem addressed in this study becomes increasingly important as genebank genomic studies increase in number, alongside other studies containing large numbers of taxa where establishing intuitive taxonomic assignments is an advantage. However, a one-size-fits-all approach is unlikely ever to arise, owing to inconsistency in a) the genetic structure of the groups being analysed, b) the conventional bases of taxonomic delineation within the taxon, and c) the taxonomic level on which differentiation is needed. Here we analysed taxa that are classified as different species, which might have a stronger evolutionary basis than, e.g., intraspecific entities like subspecies, varieties and cultivars. Expanding the small pool of current solutions is therefore advantageous, as is expanding the pool of methods used to assess their effectiveness to include more intuitive, visual methods. The solution we present here contributes to solving this problem by a) introducing a novel approach to both the classification and visual assessment steps that works well, and b) demonstrating their effectiveness on a state-of-the-art dataset that will immediately benefit a large active research community working on an agronomically and culturally important crop.

## Author Declaration

*The authors declare that the research was conducted in the absence of any commercial or financial relationships that could be construed as a potential conflict of interest*.

## Author Contributions

M.T.R-W. conceived the study, performed the analysis, and wrote the manuscript. N.S. coordinated resources and expertise and provided critical feedback on the manuscript.

## Acknowledgments

Thanks to everyone in the G2P-sol project for their encouragement and many useful and inspiring discussions, to Frank Blattner for invaluable input on the science and editing the manuscript, and to Alexandra Elbakyan and her team for indispensable support accessing resources.

## Supplementary Data

S1: Results of testing multiple parameter combinations on the classification of samples in the G2P-Sol *Capsicum* dataset using the kernel-based method presented in this study. The three panels of each row represent the results of taxonomic reclassification of the G2P pepper dataset using the kernel-based method described in the main text, under a particular parameter combination given in the title of the leftmost panel. ***Left***: The distribution of IBS distances between members of the Capsicum dataset, shown using a kernel density approximation (R v4.0.5, function `density()`) with bandwidth as indicated below each panel. The height of the blue line (arbitrarily scaled) indicates the relative weighting given to distances under that particular parameter selection. ***Centre***: The distribution of dominant proportional scores for each possible assignment for each individual, shown as per left panel. For example, if an individual receives assignment scores of 20 for taxon A, 30 for taxon B, and 50 for taxon C, then C has the dominant score, with a proportional value of 50 / (20 + 30 + 50) = 0.5. ***Right***: Taxonomic assignments of individuals in the (duplicate-purged) dataset, visualised using the t-SNE procedure with perplexity 30, as per main text figure 1. Colours as per main text fig. 1.

S2: As per S1, but showing results of k-nearest-neighbour classification with different parameter values for k.

S3: Table of assignments given to the G2P-Sol pepper accessions in this study.

## Data Availability Statement

The datasets analysed for this study are publicly available and hosted at zenodo.com under DOI:10.5281/zenodo.7016070. Scripts implementing the method in R for these data are available at github.com/mtrw/genebank_taxonomic_assignment.

