## Supplemental figures 1 for "Genebank genomics allows greatly improved taxonomic correction for *Capsicum spp*. accessions using a novel automated classification method"

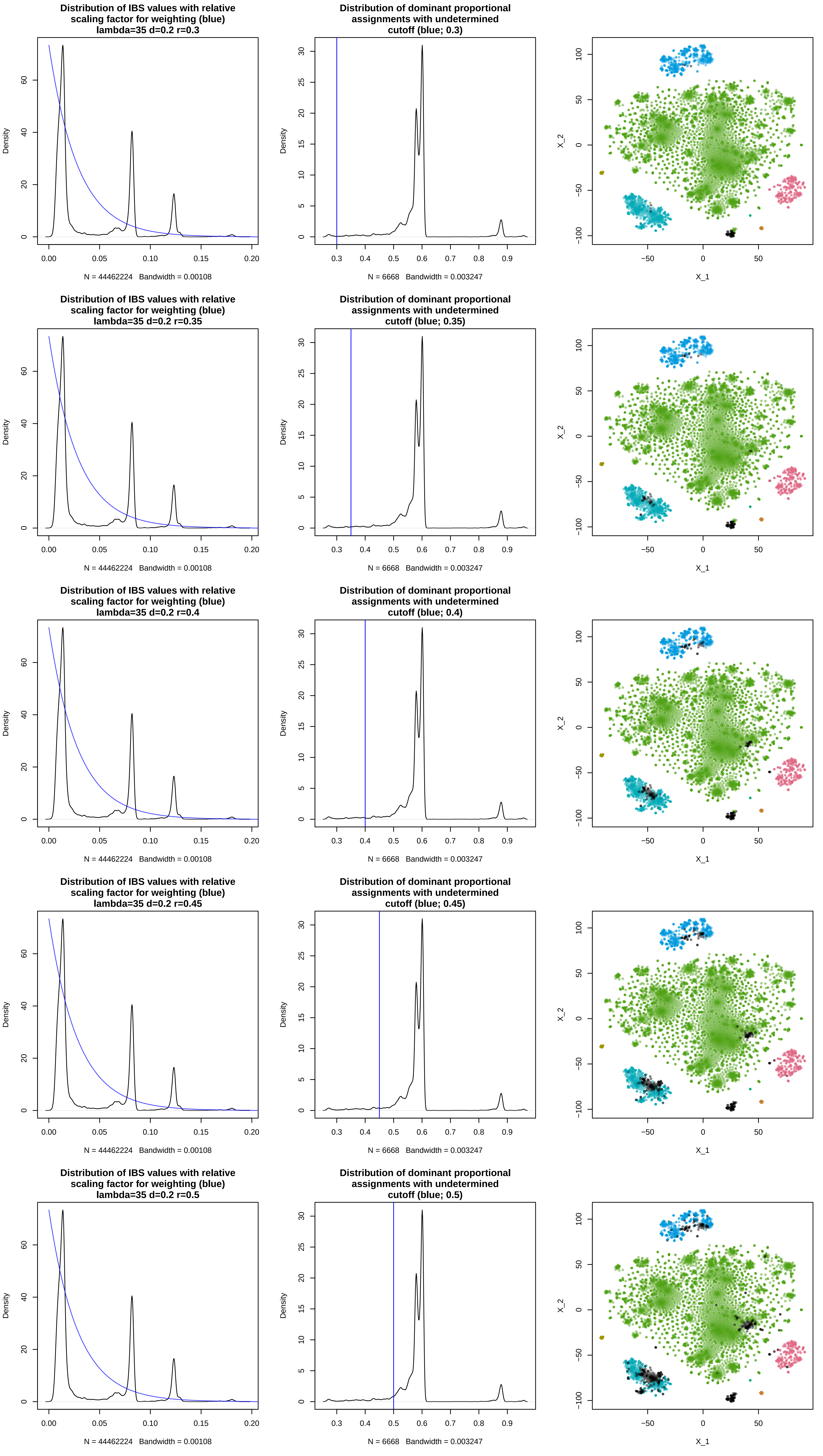
Distribution of IBS values with relative scaling factor for weighting (blue) lambda=35 d=0.2 r=0.3
Density
N = 44462224 Bandwidth = 0.00108
Distribution of dominant proportional assignments with undetermined cutoff (blue; 0.3)
Density
N = 6668 Bandwidth = 0.003247
X_1
X_2
Distribution of IBS values with relative scaling factor for weighting (blue) lambda=35 d=0.2 r=0.35
Density
N = 44462224 Bandwidth = 0.00108
Distribution of dominant proportional assignments with undetermined cutoff (blue; 0.35)
Density
N = 6668 Bandwidth = 0.003247
X_1
X_2
Distribution of IBS values with relative scaling factor for weighting (blue) lambda=35 d=0.2 r=0.4
Density
N = 44462224 Bandwidth = 0.00108
Distribution of dominant proportional assignments with undetermined cutoff (blue; 0.4)
Density
N = 6668 Bandwidth = 0.003247
X_1
X_2
Distribution of IBS values with relative scaling factor for weighting (blue) lambda=35 d=0.2 r=0.45
Density
N = 44462224 Bandwidth = 0.00108
Distribution of dominant proportional assignments with undetermined cutoff (blue; 0.45)
Density
N = 6668 Bandwidth = 0.003247
X_1
X_2
Distribution of IBS values with relative scaling factor for weighting (blue) lambda=35 d=0.2 r=0.5
Density
N = 44462224 Bandwidth = 0.00108
Distribution of dominant proportional assignments with undetermined cutoff (blue; 0.5)
Density
N = 6668 Bandwidth = 0.003247
X_1
X_2

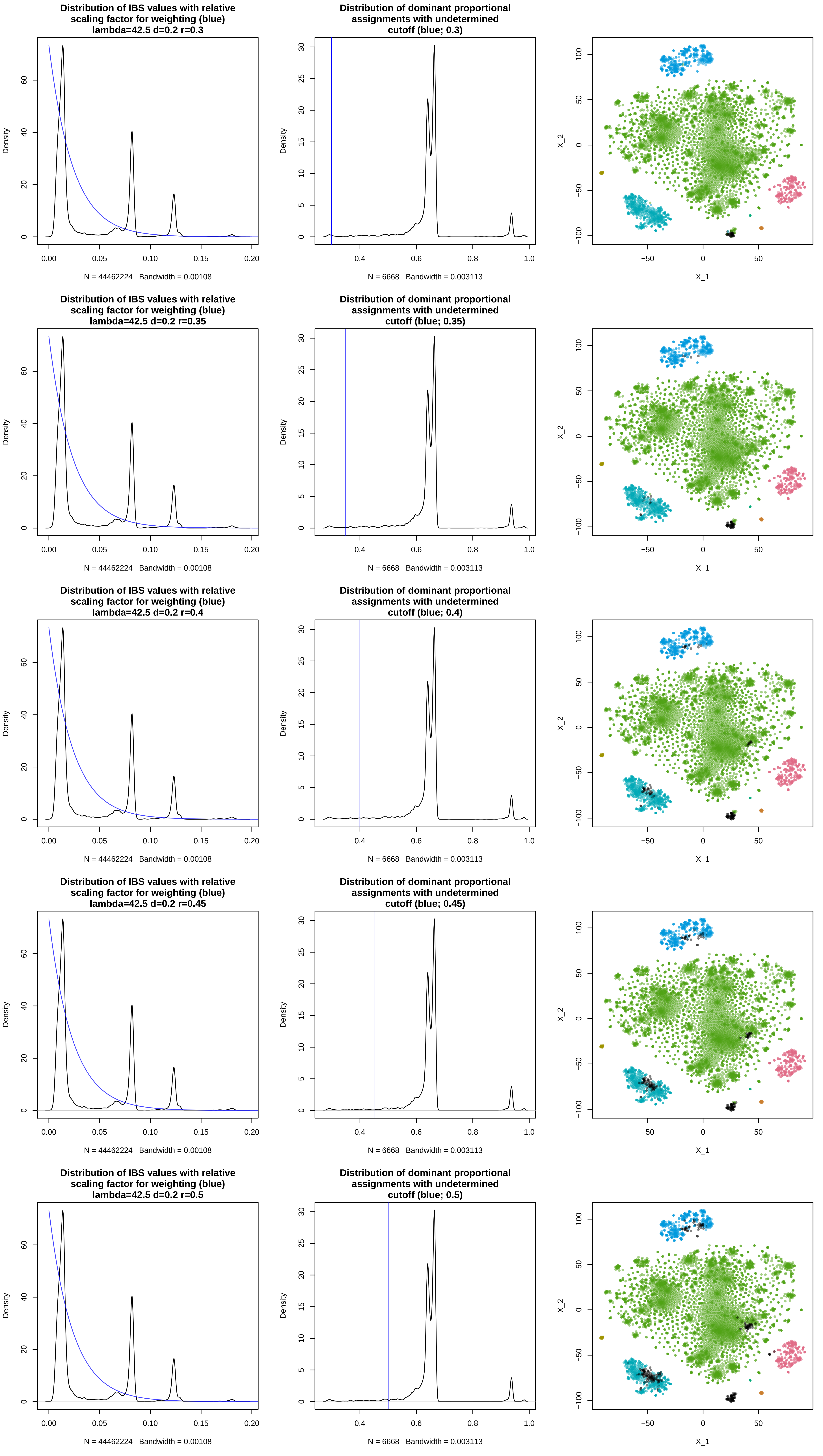
Distribution of IBS values with relative scaling factor for weighting (blue) lambda=42.5 d=0.2 r=0.3
Density
N = 44462224 Bandwidth = 0.00108
Distribution of dominant proportional assignments with undetermined cutoff (blue; 0.3)
Density
N = 6668 Bandwidth = 0.003113
X_2
X_1
Distribution of IBS values with relative scaling factor for weighting (blue) lambda=42.5 d=0.2 r=0.35
Density
N = 44462224 Bandwidth = 0.00108
Distribution of dominant proportional assignments with undetermined cutoff (blue; 0.35)
Density
N = 6668 Bandwidth = 0.003113
X_2
X_1
Distribution of IBS values with relative scaling factor for weighting (blue) lambda=42.5 d=0.2 r=0.4
Density
N = 44462224 Bandwidth = 0.00108
Distribution of dominant proportional assignments with undetermined cutoff (blue; 0.4)
Density
N = 6668 Bandwidth = 0.003113
X_2
X_1
Distribution of IBS values with relative scaling factor for weighting (blue) lambda=42.5 d=0.2 r=0.45
Density
N = 44462224 Bandwidth = 0.00108
Distribution of dominant proportional assignments with undetermined cutoff (blue; 0.45)
Density
N = 6668 Bandwidth = 0.003113
X_2
X_1
Distribution of IBS values with relative scaling factor for weighting (blue) lambda=42.5 d=0.2 r=0.5
Density
N = 44462224 Bandwidth = 0.00108
Distribution of dominant proportional assignments with undetermined cutoff (blue; 0.5)
Density
N = 6668 Bandwidth = 0.003113
X_2
X_1

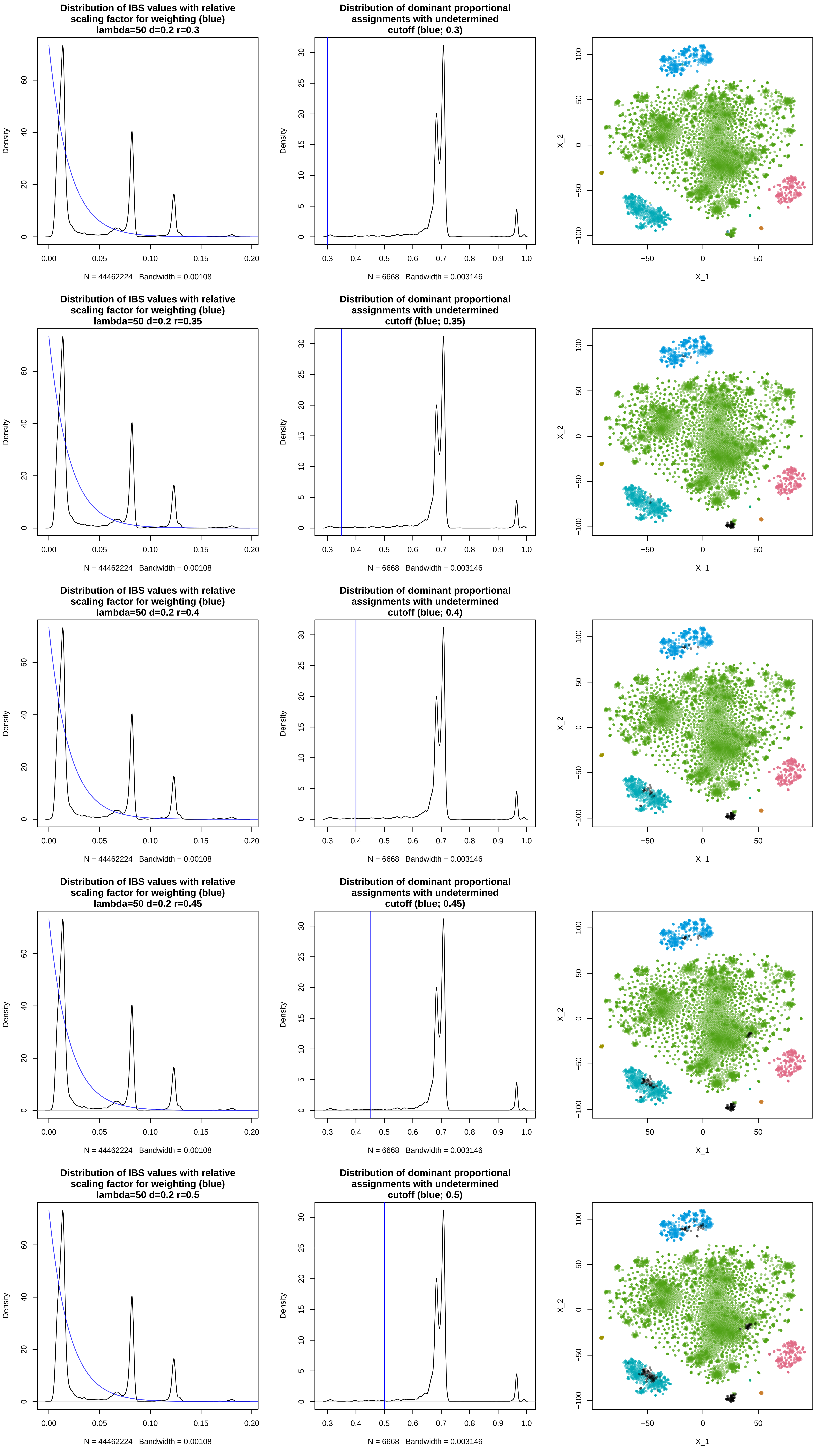
Distribution of IBS values with relative scaling factor for weighting (blue) lambda=50 d=0.2 r=0.3
Density
N = 44462224 Bandwidth = 0.00108
Distribution of dominant proportional assignments with undetermined cutoff (blue; 0.3)
Density
N = 6668 Bandwidth = 0.003146
X_2
X_1
Distribution of IBS values with relative scaling factor for weighting (blue) lambda=50 d=0.2 r=0.35
Density
N = 44462224 Bandwidth = 0.00108
Distribution of dominant proportional assignments with undetermined cutoff (blue; 0.35)
Density
N = 6668 Bandwidth = 0.003146
X_2
X_1
Distribution of IBS values with relative scaling factor for weighting (blue) lambda=50 d=0.2 r=0.4
Density
N = 44462224 Bandwidth = 0.00108
Distribution of dominant proportional assignments with undetermined cutoff (blue; 0.4)
Density
N = 6668 Bandwidth = 0.003146
X_2
X_1
Distribution of IBS values with relative scaling factor for weighting (blue) lambda=50 d=0.2 r=0.45
Density
N = 44462224 Bandwidth = 0.00108
Distribution of dominant proportional assignments with undetermined cutoff (blue; 0.45)
Density
N = 6668 Bandwidth = 0.003146
X_2
X_1
Distribution of IBS values with relative scaling factor for weighting (blue) lambda=50 d=0.2 r=0.5
Density
N = 44462224 Bandwidth = 0.00108
Distribution of dominant proportional assignments with undetermined cutoff (blue; 0.5)
Density
N = 6668 Bandwidth = 0.003146
X_2
X_1

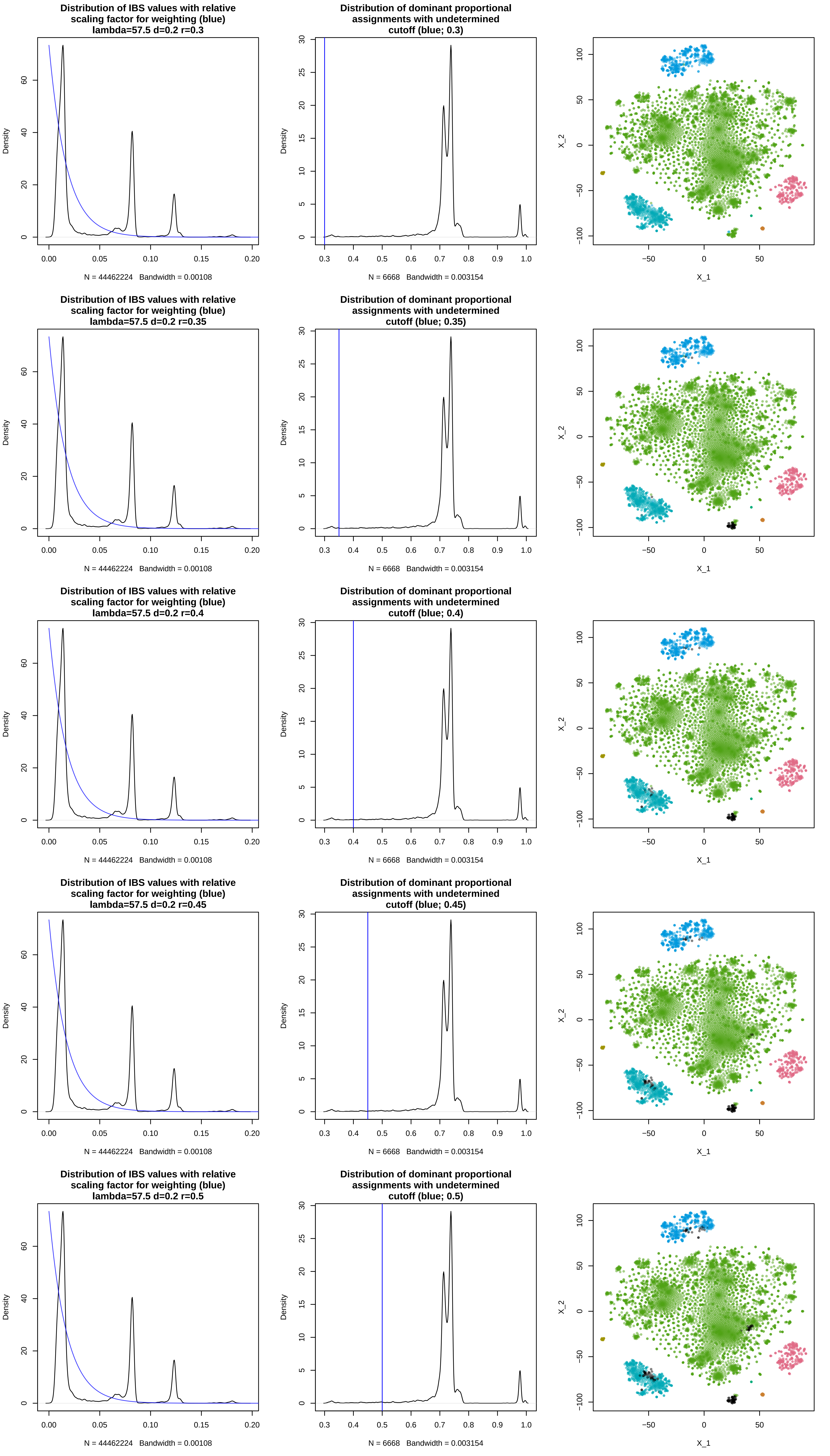

Distribution of IBS values with relative scaling factor for weighting (blue) lambda=57.5 d=0.2 r=0.3
Density
N = 44462224 Bandwidth = 0.00108
Distribution of dominant proportional assignments with undetermined cutoff (blue; 0.3)
Density
N = 6668 Bandwidth = 0.003154
X_1
X_2
Distribution of IBS values with relative scaling factor for weighting (blue) lambda=57.5 d=0.2 r=0.35
Density
N = 44462224 Bandwidth = 0.00108
Distribution of dominant proportional assignments with undetermined cutoff (blue; 0.35)
Density
N = 6668 Bandwidth = 0.003154
X_1
X_2
Distribution of IBS values with relative scaling factor for weighting (blue) lambda=57.5 d=0.2 r=0.4
Density
N = 44462224 Bandwidth = 0.00108
Distribution of dominant proportional assignments with undetermined cutoff (blue; 0.4)
Density
N = 6668 Bandwidth = 0.003154
X_1
X_2
Distribution of IBS values with relative scaling factor for weighting (blue) lambda=57.5 d=0.2 r=0.45
Density
N = 44462224 Bandwidth = 0.00108
Distribution of dominant proportional assignments with undetermined cutoff (blue; 0.45)
Density
N = 6668 Bandwidth = 0.003154
X_1
X_2
Distribution of IBS values with relative scaling factor for weighting (blue) lambda=57.5 d=0.2 r=0.5
Density
N = 44462224 Bandwidth = 0.00108
Distribution of dominant proportional assignments with undetermined cutoff (blue; 0.5)
Density
N = 6668 Bandwidth = 0.003154
X_1
X_2

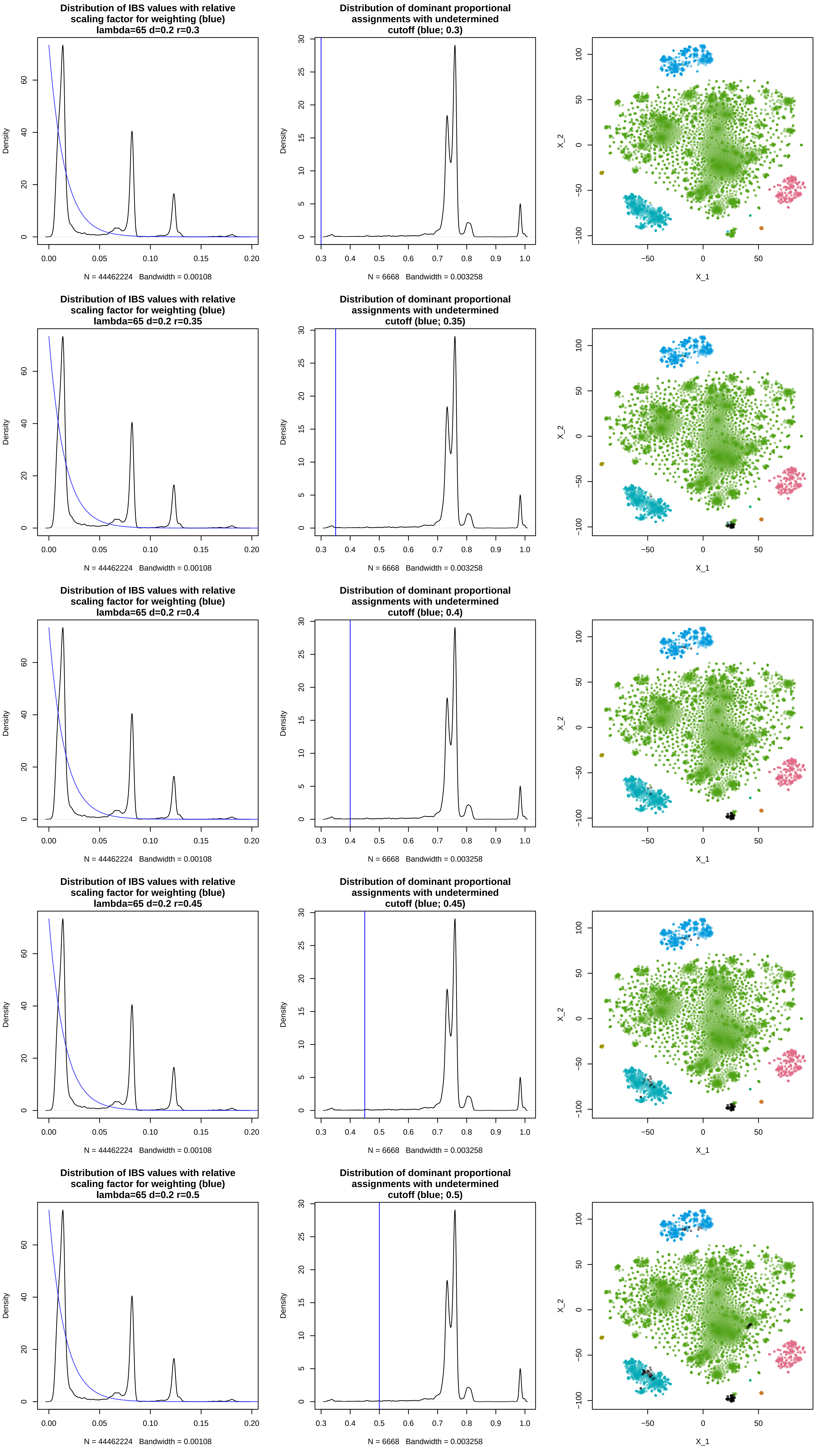
Distribution of IBS values with relative scaling factor for weighting (blue) lambda=65 d=0.2 r=0.3
Density
N = 44462224 Bandwidth = 0.00108
Distribution of dominant proportional assignments with undetermined cutoff (blue; 0.3)
Density
N = 6668 Bandwidth = 0.003258
X_2
X_1
Distribution of IBS values with relative scaling factor for weighting (blue) lambda=65 d=0.2 r=0.35
Density
N = 44462224 Bandwidth = 0.00108
Distribution of dominant proportional assignments with undetermined cutoff (blue; 0.35)
Density
N = 6668 Bandwidth = 0.003258
X_2
X_1
Distribution of IBS values with relative scaling factor for weighting (blue) lambda=65 d=0.2 r=0.4
Density
N = 44462224 Bandwidth = 0.00108
Distribution of dominant proportional assignments with undetermined cutoff (blue; 0.4)
Density
N = 6668 Bandwidth = 0.003258
X_2
X_1
Distribution of IBS values with relative scaling factor for weighting (blue) lambda=65 d=0.2 r=0.45
Density
N = 44462224 Bandwidth = 0.00108
Distribution of dominant proportional assignments with undetermined cutoff (blue; 0.45)
Density
N = 6668 Bandwidth = 0.003258
X_2
X_1
Distribution of IBS values with relative scaling factor for weighting (blue) lambda=65 d=0.2 r=0.5
Density
N = 44462224 Bandwidth = 0.00108
Distribution of dominant proportional assignments with undetermined cutoff (blue; 0.5)
Density
N = 6668 Bandwidth = 0.003258
X_2
X_1

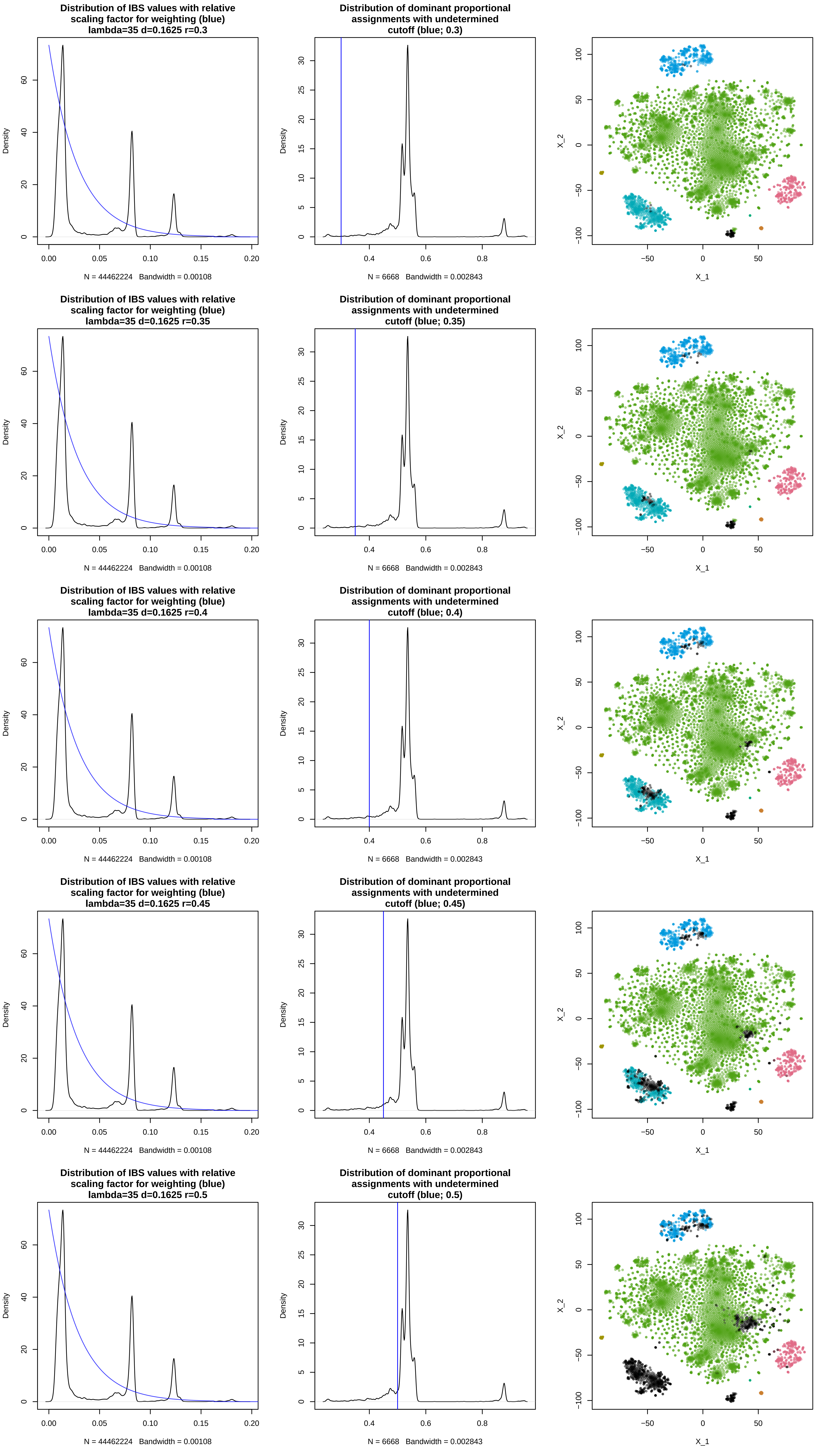

Distribution of IBS values with relative scaling factor for weighting (blue) lambda=35 d=0.1625 r=0.3
Distribution of dominant proportional assignments with undetermined cutoff (blue; 0.3)
Distribution of IBS values with relative scaling factor for weighting (blue) lambda=35 d=0.1625 r=0.35
Distribution of dominant proportional assignments with undetermined cutoff (blue; 0.35)
Distribution of IBS values with relative scaling factor for weighting (blue) lambda=35 d=0.1625 r=0.4
Distribution of dominant proportional assignments with undetermined cutoff (blue; 0.4)
Distribution of IBS values with relative scaling factor for weighting (blue) lambda=35 d=0.1625 r=0.45
Distribution of dominant proportional assignments with undetermined cutoff (blue; 0.45)
Distribution of IBS values with relative scaling factor for weighting (blue) lambda=35 d=0.1625 r=0.5
Distribution of dominant proportional assignments with undetermined cutoff (blue; 0.5)
Density
X_1
X_2
N = 44462224 Bandwidth = 0.00108
N = 6668 Bandwidth = 0.002843

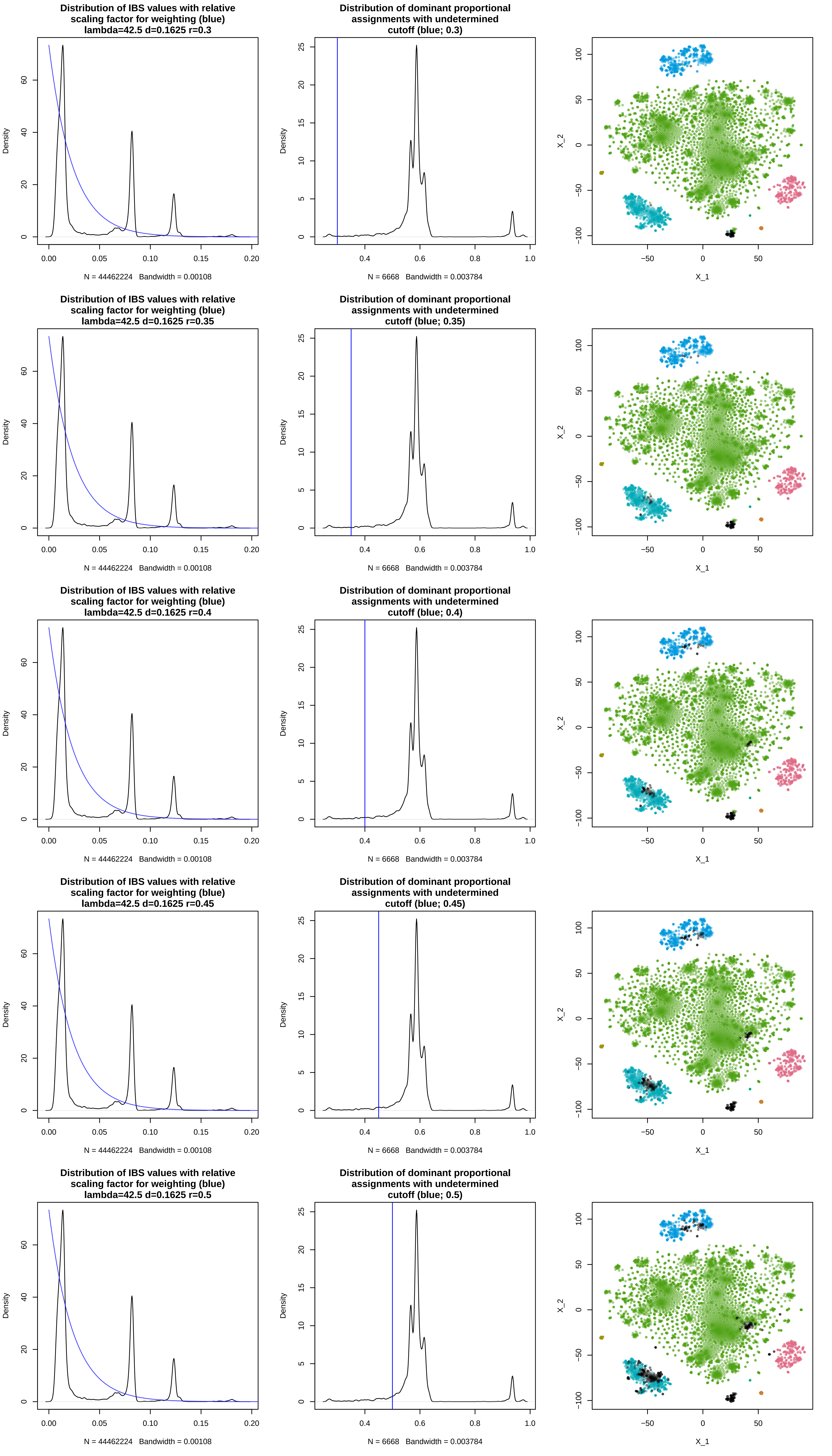
Distribution of IBS values with relative scaling factor for weighting (blue) lambda=42.5 d=0.1625 r=0.3
Density
N = 44462224 Bandwidth = 0.00108
Distribution of dominant proportional assignments with undetermined cutoff (blue; 0.3)
Density
N = 6668 Bandwidth = 0.003784
X_1
X_2
Distribution of IBS values with relative scaling factor for weighting (blue) lambda=42.5 d=0.1625 r=0.35
Density
N = 44462224 Bandwidth = 0.00108
Distribution of dominant proportional assignments with undetermined cutoff (blue; 0.35)
Density
N = 6668 Bandwidth = 0.003784
X_1
X_2
Distribution of IBS values with relative scaling factor for weighting (blue) lambda=42.5 d=0.1625 r=0.4
Density
N = 44462224 Bandwidth = 0.00108
Distribution of dominant proportional assignments with undetermined cutoff (blue; 0.4)
Density
N = 6668 Bandwidth = 0.003784
X_1
X_2
Distribution of IBS values with relative scaling factor for weighting (blue) lambda=42.5 d=0.1625 r=0.45
Density
N = 44462224 Bandwidth = 0.00108
Distribution of dominant proportional assignments with undetermined cutoff (blue; 0.45)
Density
N = 6668 Bandwidth = 0.003784
X_1
X_2
Distribution of IBS values with relative scaling factor for weighting (blue) lambda=42.5 d=0.1625 r=0.5
Density
N = 44462224 Bandwidth = 0.00108
Distribution of dominant proportional assignments with undetermined cutoff (blue; 0.5)
Density
N = 6668 Bandwidth = 0.003784
X_1
X_2

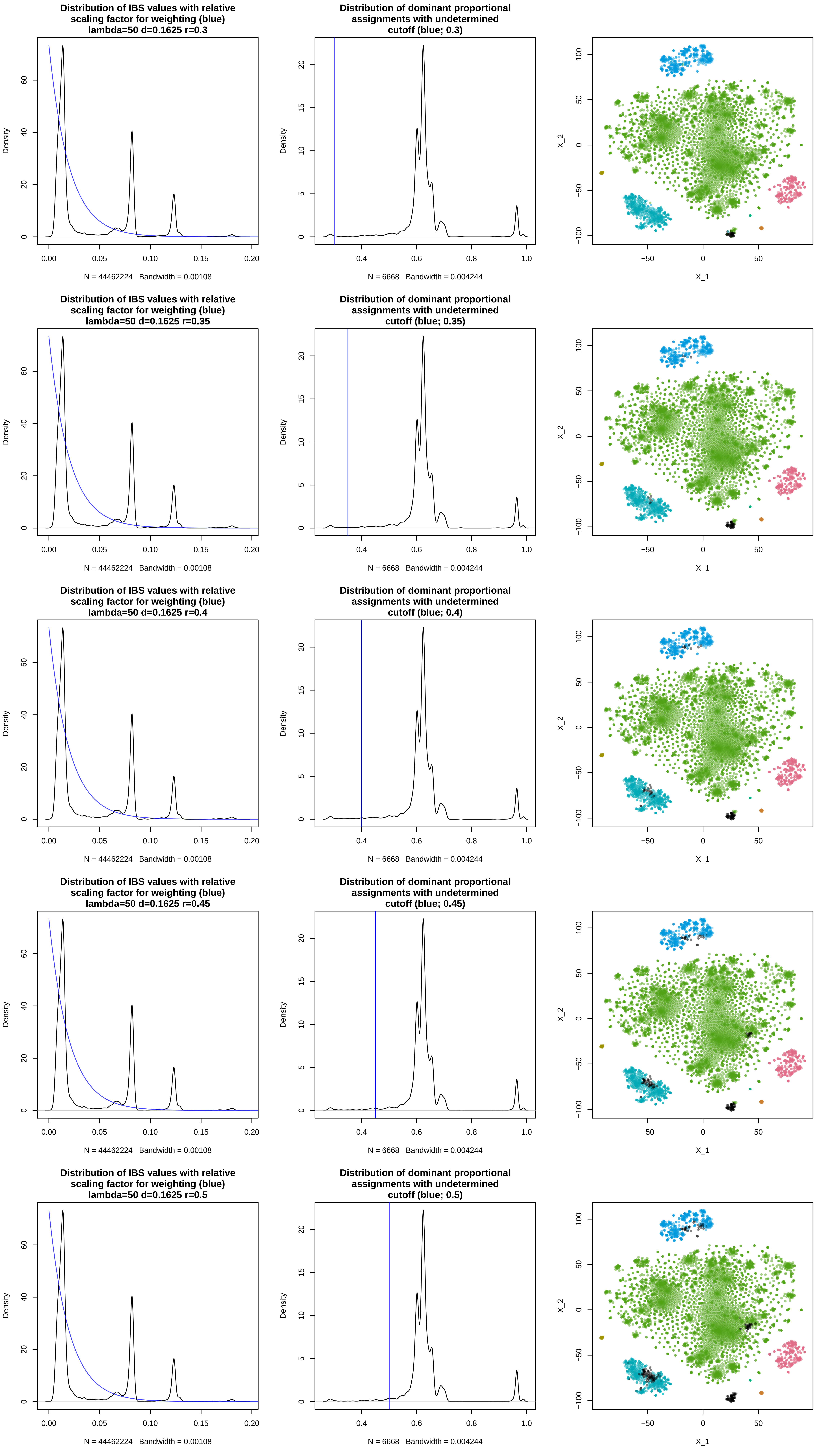

Distribution of IBS values with relative scaling factor for weighting (blue) lambda=50 d=0.1625 r=0.3
Density
N = 44462224 Bandwidth = 0.00108
Distribution of dominant proportional assignments with undetermined cutoff (blue; 0.3)
Density
N = 6668 Bandwidth = 0.004244
X_2
X_1
Distribution of IBS values with relative scaling factor for weighting (blue) lambda=50 d=0.1625 r=0.35
Density
N = 44462224 Bandwidth = 0.00108
Distribution of dominant proportional assignments with undetermined cutoff (blue; 0.35)
Density
N = 6668 Bandwidth = 0.004244
X_2
X_1
Distribution of IBS values with relative scaling factor for weighting (blue) lambda=50 d=0.1625 r=0.4
Density
N = 44462224 Bandwidth = 0.00108
Distribution of dominant proportional assignments with undetermined cutoff (blue; 0.4)
Density
N = 6668 Bandwidth = 0.004244
X_2
X_1
Distribution of IBS values with relative scaling factor for weighting (blue) lambda=50 d=0.1625 r=0.45
Density
N = 44462224 Bandwidth = 0.00108
Distribution of dominant proportional assignments with undetermined cutoff (blue; 0.45)
Density
N = 6668 Bandwidth = 0.004244
X_2
X_1
Distribution of IBS values with relative scaling factor for weighting (blue) lambda=50 d=0.1625 r=0.5
Density
N = 44462224 Bandwidth = 0.00108
Distribution of dominant proportional assignments with undetermined cutoff (blue; 0.5)
Density
N = 6668 Bandwidth = 0.004244
X_2
X_1

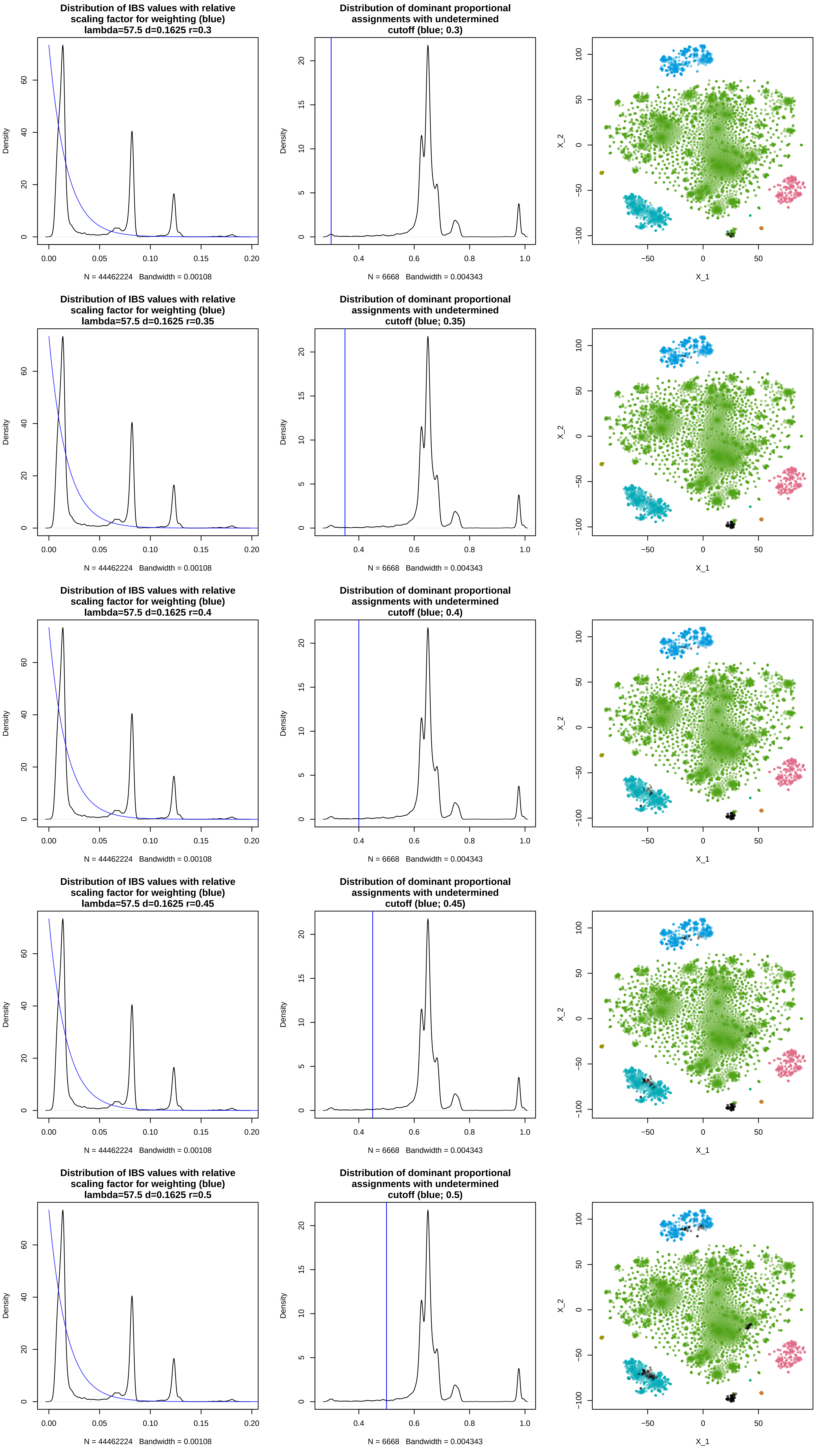

Distribution of IBS values with relative scaling factor for weighting (blue) lambda=57.5 d=0.1625 r=0.3
Distribution of dominant proportional assignments with undetermined cutoff (blue; 0.3)
Distribution of IBS values with relative scaling factor for weighting (blue) lambda=57.5 d=0.1625 r=0.35
Distribution of dominant proportional assignments with undetermined cutoff (blue; 0.35)
Distribution of IBS values with relative scaling factor for weighting (blue) lambda=57.5 d=0.1625 r=0.4
Distribution of dominant proportional assignments with undetermined cutoff (blue; 0.4)
Distribution of IBS values with relative scaling factor for weighting (blue) lambda=57.5 d=0.1625 r=0.45
Distribution of dominant proportional assignments with undetermined cutoff (blue; 0.45)
Distribution of IBS values with relative scaling factor for weighting (blue) lambda=57.5 d=0.1625 r=0.5
Distribution of dominant proportional assignments with undetermined cutoff (blue; 0.5)
Density
N = 44462224 Bandwidth = 0.00108
N = 6668 Bandwidth = 0.004343
X_1
X_2

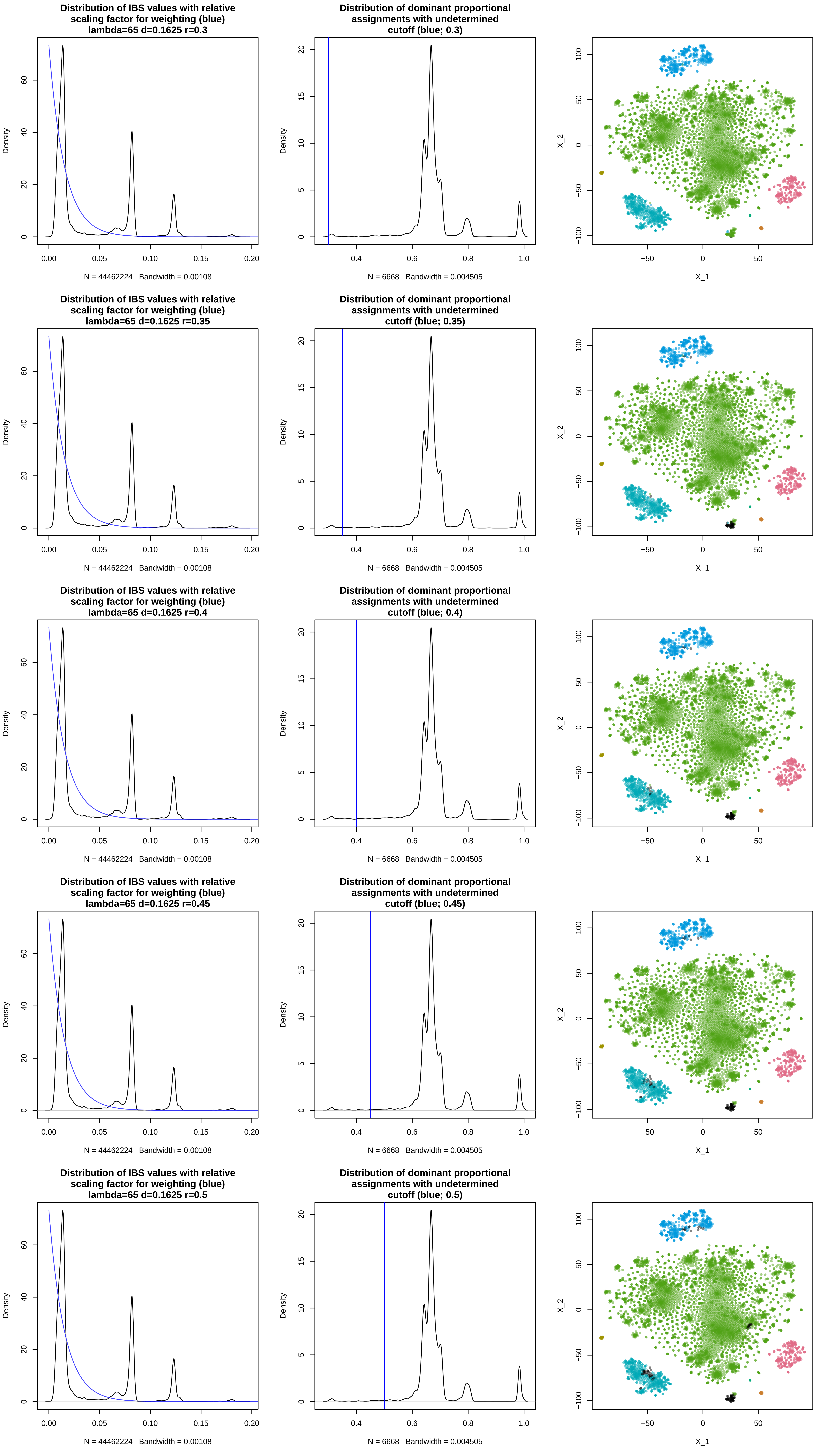

Distribution of IBS values with relative scaling factor for weighting (blue) lambda=65 d=0.1625 r=0.3
Density
N = 44462224 Bandwidth = 0.00108
Distribution of dominant proportional assignments with undetermined cutoff (blue; 0.3)
Density
N = 6668 Bandwidth = 0.004505
X_2
X_1
Distribution of IBS values with relative scaling factor for weighting (blue) lambda=65 d=0.1625 r=0.35
Density
N = 44462224 Bandwidth = 0.00108
Distribution of dominant proportional assignments with undetermined cutoff (blue; 0.35)
Density
N = 6668 Bandwidth = 0.004505
X_2
X_1
Distribution of IBS values with relative scaling factor for weighting (blue) lambda=65 d=0.1625 r=0.4
Density
N = 44462224 Bandwidth = 0.00108
Distribution of dominant proportional assignments with undetermined cutoff (blue; 0.4)
Density
N = 6668 Bandwidth = 0.004505
X_2
X_1
Distribution of IBS values with relative scaling factor for weighting (blue) lambda=65 d=0.1625 r=0.45
Density
N = 44462224 Bandwidth = 0.00108
Distribution of dominant proportional assignments with undetermined cutoff (blue; 0.45)
Density
N = 6668 Bandwidth = 0.004505
X_2
X_1
Distribution of IBS values with relative scaling factor for weighting (blue) lambda=65 d=0.1625 r=0.5
Density
N = 44462224 Bandwidth = 0.00108
Distribution of dominant proportional assignments with undetermined cutoff (blue; 0.5)
Density
N = 6668 Bandwidth = 0.004505
X_2
X_1

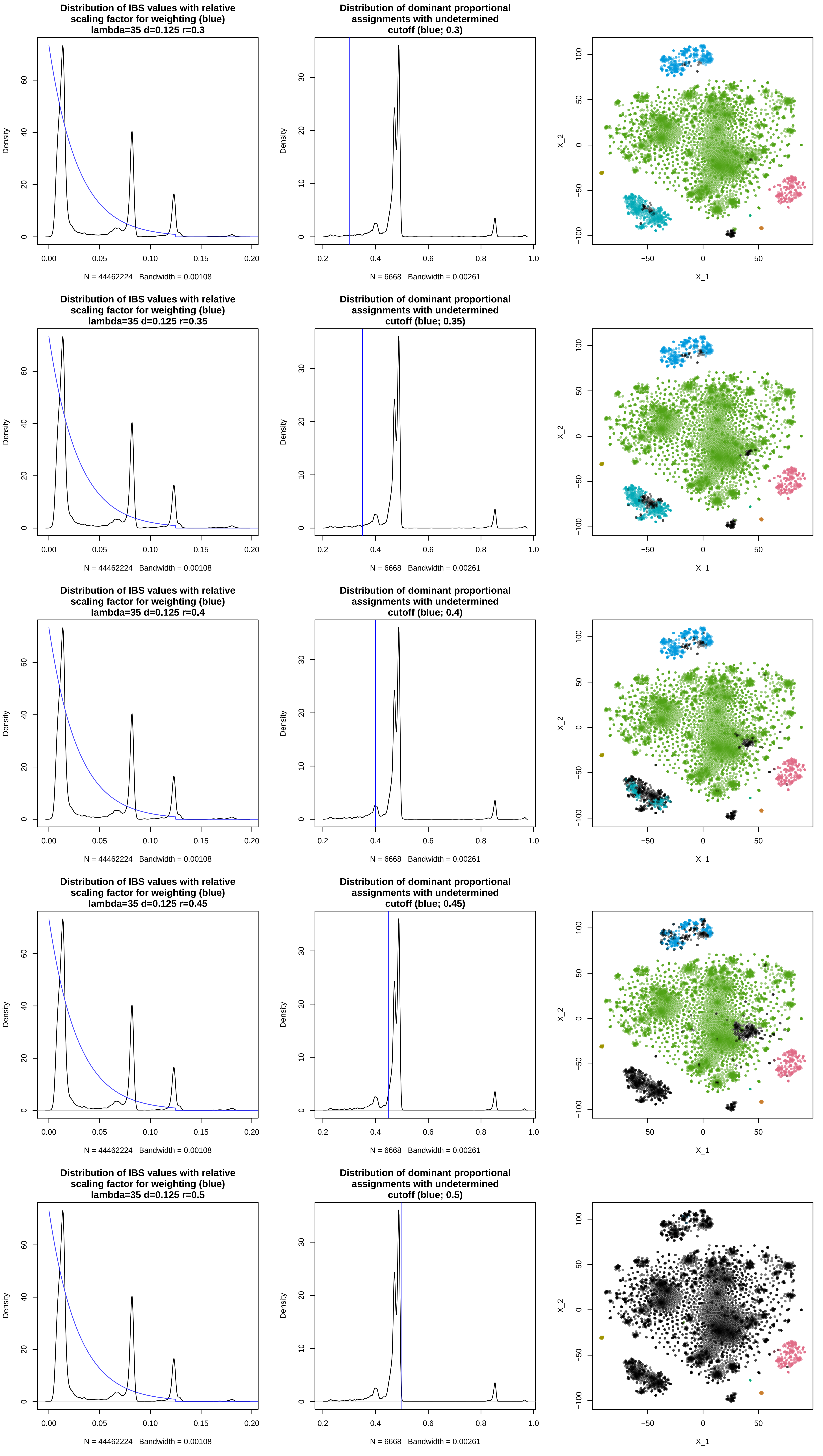
Distribution of IBS values with relative scaling factor for weighting (blue) lambda=35 d=0.125 r=0.3
Density
N = 44462224 Bandwidth = 0.00108
Distribution of dominant proportional assignments with undetermined cutoff (blue; 0.3)
Density
N = 6668 Bandwidth = 0.00261
X_1
X_2
Distribution of IBS values with relative scaling factor for weighting (blue) lambda=35 d=0.125 r=0.35
Density
N = 44462224 Bandwidth = 0.00108
Distribution of dominant proportional assignments with undetermined cutoff (blue; 0.35)
Density
N = 6668 Bandwidth = 0.00261
X_1
X_2
Distribution of IBS values with relative scaling factor for weighting (blue) lambda=35 d=0.125 r=0.4
Density
N = 44462224 Bandwidth = 0.00108
Distribution of dominant proportional assignments with undetermined cutoff (blue; 0.4)
Density
N = 6668 Bandwidth = 0.00261
X_1
X_2
Distribution of IBS values with relative scaling factor for weighting (blue) lambda=35 d=0.125 r=0.45
Density
N = 44462224 Bandwidth = 0.00108
Distribution of dominant proportional assignments with undetermined cutoff (blue; 0.45)
Density
N = 6668 Bandwidth = 0.00261
X_1
X_2
Distribution of IBS values with relative scaling factor for weighting (blue) lambda=35 d=0.125 r=0.5
Density
N = 44462224 Bandwidth = 0.00108
Distribution of dominant proportional assignments with undetermined cutoff (blue; 0.5)
Density
N = 6668 Bandwidth = 0.00261
X_1
X_2

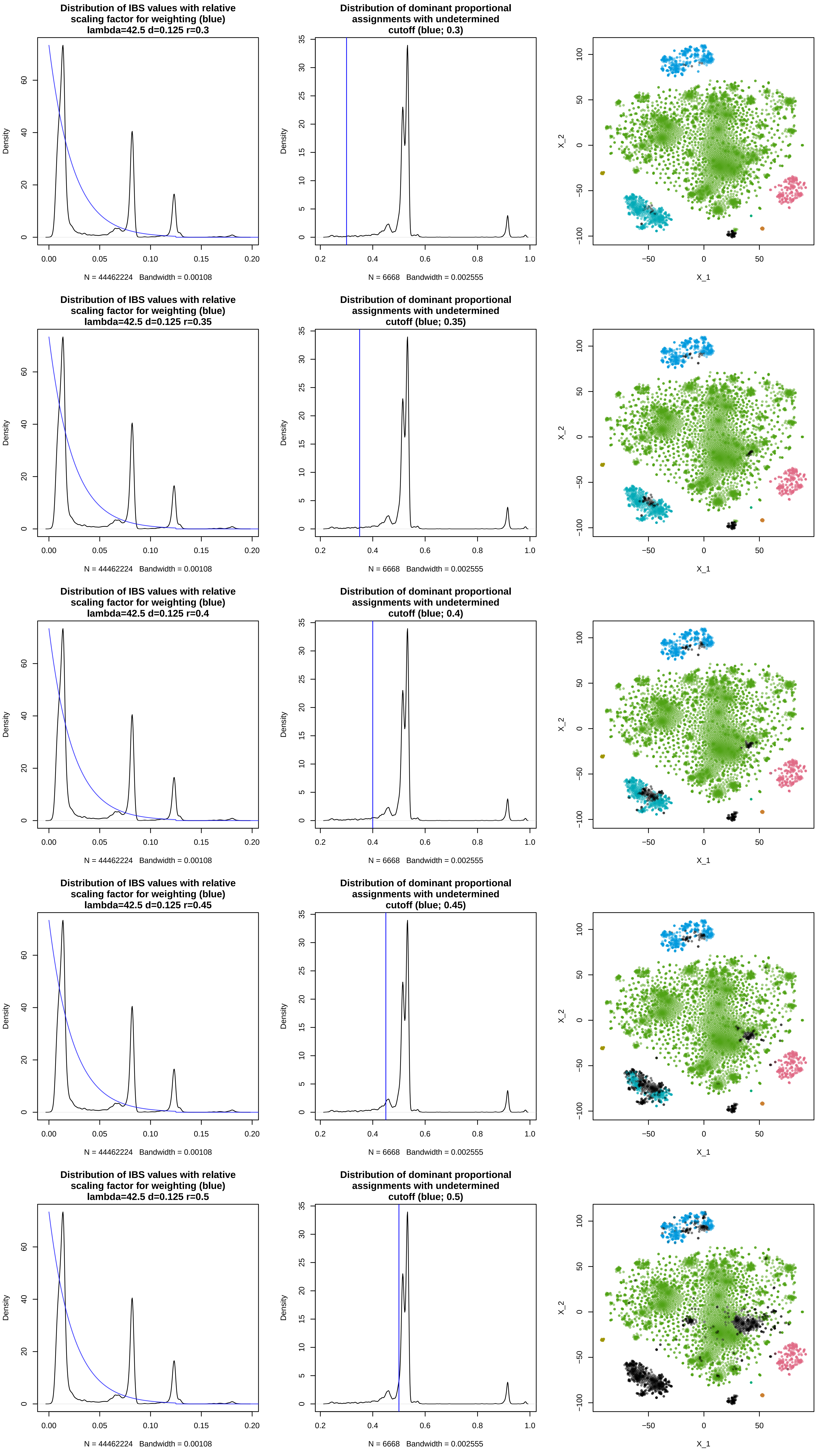
Distribution of IBS values with relative scaling factor for weighting (blue) lambda=42.5 d=0.125 r=0.3
Density
0 20 40 60
0.00 0.05 0.10 0.15 0.20
N = 44462224 Bandwidth = 0.00108
Distribution of dominant proportional assignments with undetermined cutoff (blue; 0.3)
Density
0 5 10 15 20 25 30 35
0.2 0.4 0.6 0.8 1.0
N = 6668 Bandwidth = 0.002555
X_2
−100 −50 0 50 100
X_1
−50 0 50
Distribution of IBS values with relative scaling factor for weighting (blue) lambda=42.5 d=0.125 r=0.35
N = 44462224 Bandwidth = 0.00108
Distribution of dominant proportional assignments with undetermined cutoff (blue; 0.35)
N = 6668 Bandwidth = 0.002555
Distribution of IBS values with relative scaling factor for weighting (blue) lambda=42.5 d=0.125 r=0.4
N = 44462224 Bandwidth = 0.00108
Distribution of dominant proportional assignments with undetermined cutoff (blue; 0.4)
N = 6668 Bandwidth = 0.002555
Distribution of IBS values with relative scaling factor for weighting (blue) lambda=42.5 d=0.125 r=0.45
N = 44462224 Bandwidth = 0.00108
Distribution of dominant proportional assignments with undetermined cutoff (blue; 0.45)
N = 6668 Bandwidth = 0.002555
Distribution of IBS values with relative scaling factor for weighting (blue) lambda=42.5 d=0.125 r=0.5
N = 44462224 Bandwidth = 0.00108
Distribution of dominant proportional assignments with undetermined cutoff (blue; 0.5)
N = 6668 Bandwidth = 0.002555

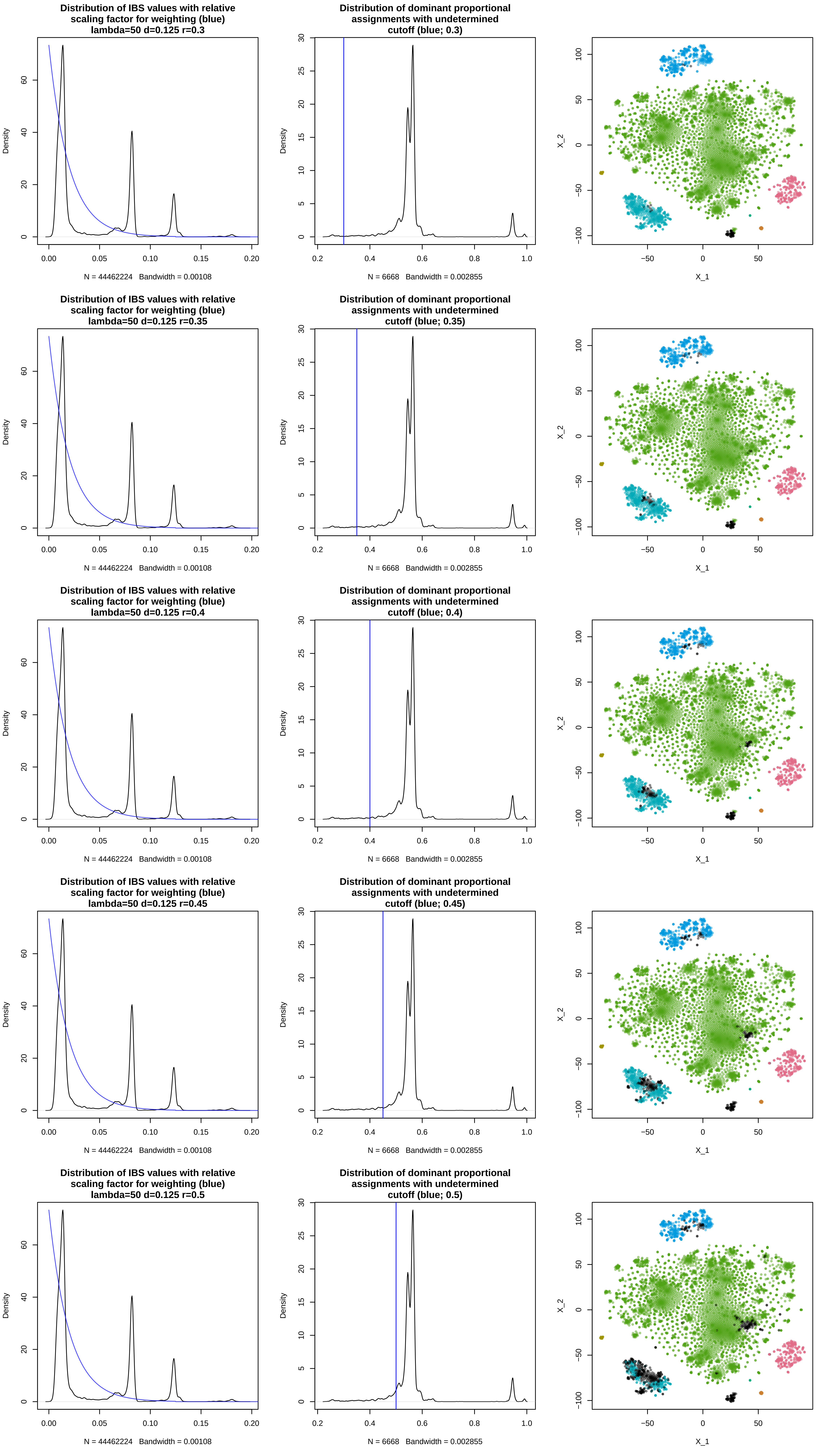
Distribution of IBS values with relative scaling factor for weighting (blue) lambda=50 d=0.125 r=0.3
Density
N = 44462224 Bandwidth = 0.00108
Distribution of dominant proportional assignments with undetermined cutoff (blue; 0.3)
Density
N = 6668 Bandwidth = 0.002855
X_1
X_2
Distribution of IBS values with relative scaling factor for weighting (blue) lambda=50 d=0.125 r=0.35
Density
N = 44462224 Bandwidth = 0.00108
Distribution of dominant proportional assignments with undetermined cutoff (blue; 0.35)
Density
N = 6668 Bandwidth = 0.002855
X_1
X_2
Distribution of IBS values with relative scaling factor for weighting (blue) lambda=50 d=0.125 r=0.4
Density
N = 44462224 Bandwidth = 0.00108
Distribution of dominant proportional assignments with undetermined cutoff (blue; 0.4)
Density
N = 6668 Bandwidth = 0.002855
X_1
X_2
Distribution of IBS values with relative scaling factor for weighting (blue) lambda=50 d=0.125 r=0.45
Density
N = 44462224 Bandwidth = 0.00108
Distribution of dominant proportional assignments with undetermined cutoff (blue; 0.45)
Density
N = 6668 Bandwidth = 0.002855
X_1
X_2
Distribution of IBS values with relative scaling factor for weighting (blue) lambda=50 d=0.125 r=0.5
Density
N = 44462224 Bandwidth = 0.00108
Distribution of dominant proportional assignments with undetermined cutoff (blue; 0.5)
Density
N = 6668 Bandwidth = 0.002855
X_1
X_2

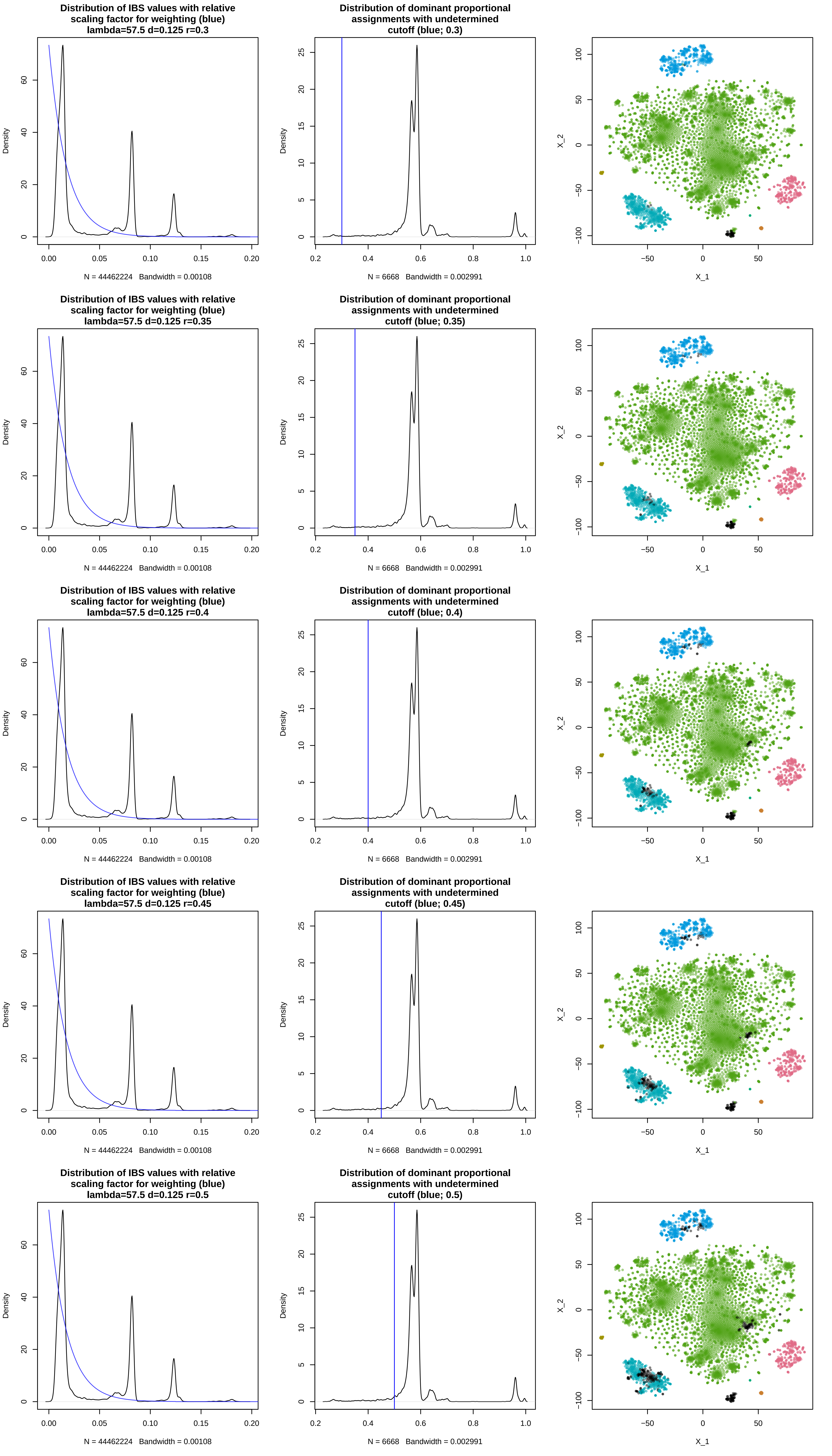

Distribution of IBS values with relative scaling factor for weighting (blue) lambda=57.5 d=0.125 r=0.3
Density
N = 44462224 Bandwidth = 0.00108
Distribution of dominant proportional assignments with undetermined cutoff (blue; 0.3)
Density
N = 6668 Bandwidth = 0.002991
X_2
X_1
Distribution of IBS values with relative scaling factor for weighting (blue) lambda=57.5 d=0.125 r=0.35
Density
N = 44462224 Bandwidth = 0.00108
Distribution of dominant proportional assignments with undetermined cutoff (blue; 0.35)
Density
N = 6668 Bandwidth = 0.002991
X_2
X_1
Distribution of IBS values with relative scaling factor for weighting (blue) lambda=57.5 d=0.125 r=0.4
Density
N = 44462224 Bandwidth = 0.00108
Distribution of dominant proportional assignments with undetermined cutoff (blue; 0.4)
Density
N = 6668 Bandwidth = 0.002991
X_2
X_1
Distribution of IBS values with relative scaling factor for weighting (blue) lambda=57.5 d=0.125 r=0.45
Density
N = 44462224 Bandwidth = 0.00108
Distribution of dominant proportional assignments with undetermined cutoff (blue; 0.45)
Density
N = 6668 Bandwidth = 0.002991
X_2
X_1
Distribution of IBS values with relative scaling factor for weighting (blue) lambda=57.5 d=0.125 r=0.5
Density
N = 44462224 Bandwidth = 0.00108
Distribution of dominant proportional assignments with undetermined cutoff (blue; 0.5)
Density
N = 6668 Bandwidth = 0.002991
X_2
X_1

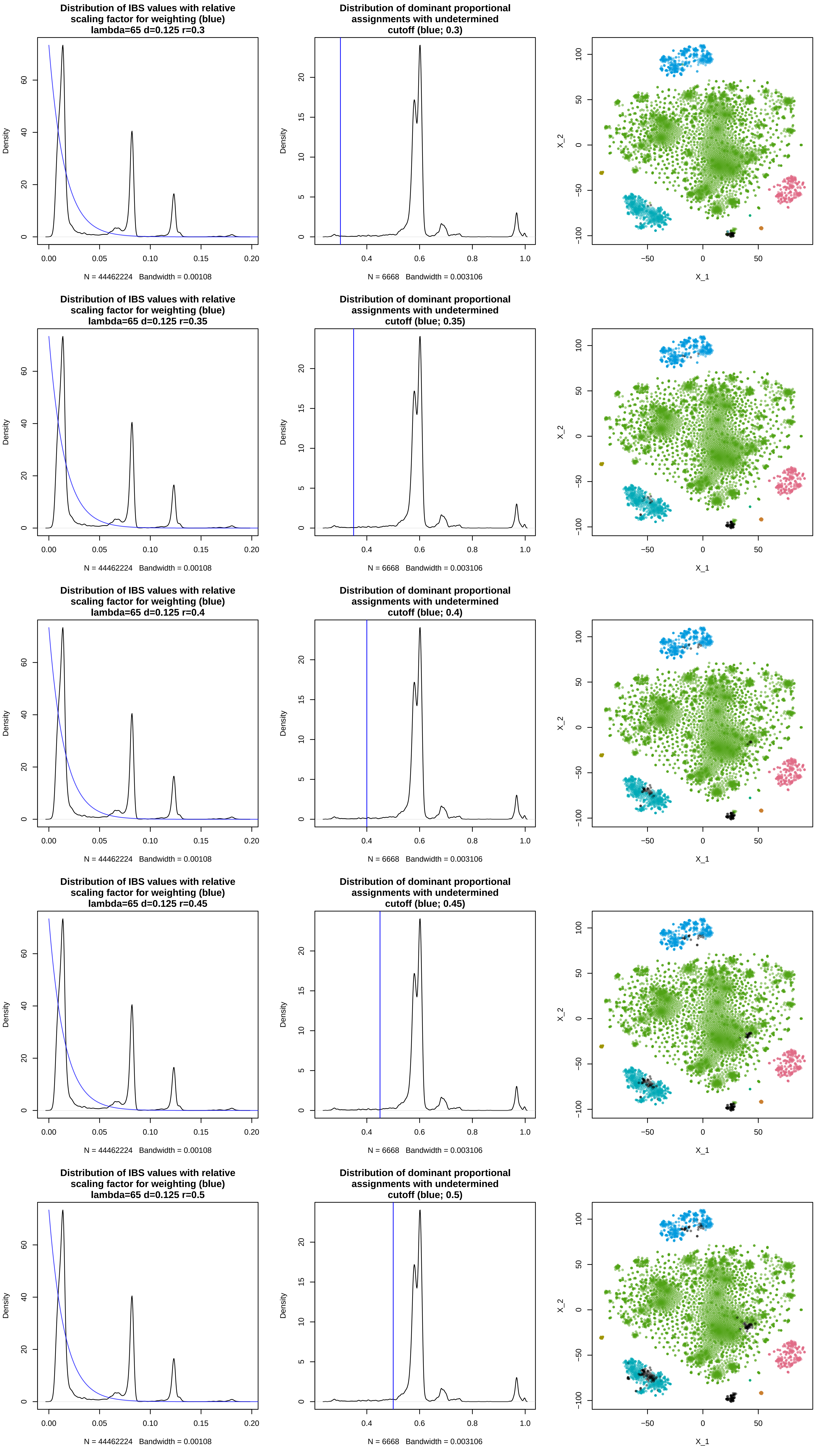
Distribution of IBS values with relative scaling factor for weighting (blue) lambda=65 d=0.125 r=0.3
Density
N = 44462224 Bandwidth = 0.00108
Distribution of dominant proportional assignments with undetermined cutoff (blue; 0.3)
Density
N = 6668 Bandwidth = 0.003106
X_2
X_1
Distribution of IBS values with relative scaling factor for weighting (blue) lambda=65 d=0.125 r=0.35
Density
N = 44462224 Bandwidth = 0.00108
Distribution of dominant proportional assignments with undetermined cutoff (blue; 0.35)
Density
N = 6668 Bandwidth = 0.003106
X_2
X_1
Distribution of IBS values with relative scaling factor for weighting (blue) lambda=65 d=0.125 r=0.4
Density
N = 44462224 Bandwidth = 0.00108
Distribution of dominant proportional assignments with undetermined cutoff (blue; 0.4)
Density
N = 6668 Bandwidth = 0.003106
X_2
X_1
Distribution of IBS values with relative scaling factor for weighting (blue) lambda=65 d=0.125 r=0.45
Density
N = 44462224 Bandwidth = 0.00108
Distribution of dominant proportional assignments with undetermined cutoff (blue; 0.45)
Density
N = 6668 Bandwidth = 0.003106
X_2
X_1
Distribution of IBS values with relative scaling factor for weighting (blue) lambda=65 d=0.125 r=0.5
Density
N = 44462224 Bandwidth = 0.00108
Distribution of dominant proportional assignments with undetermined cutoff (blue; 0.5)
Density
N = 6668 Bandwidth = 0.003106
X_2
X_1

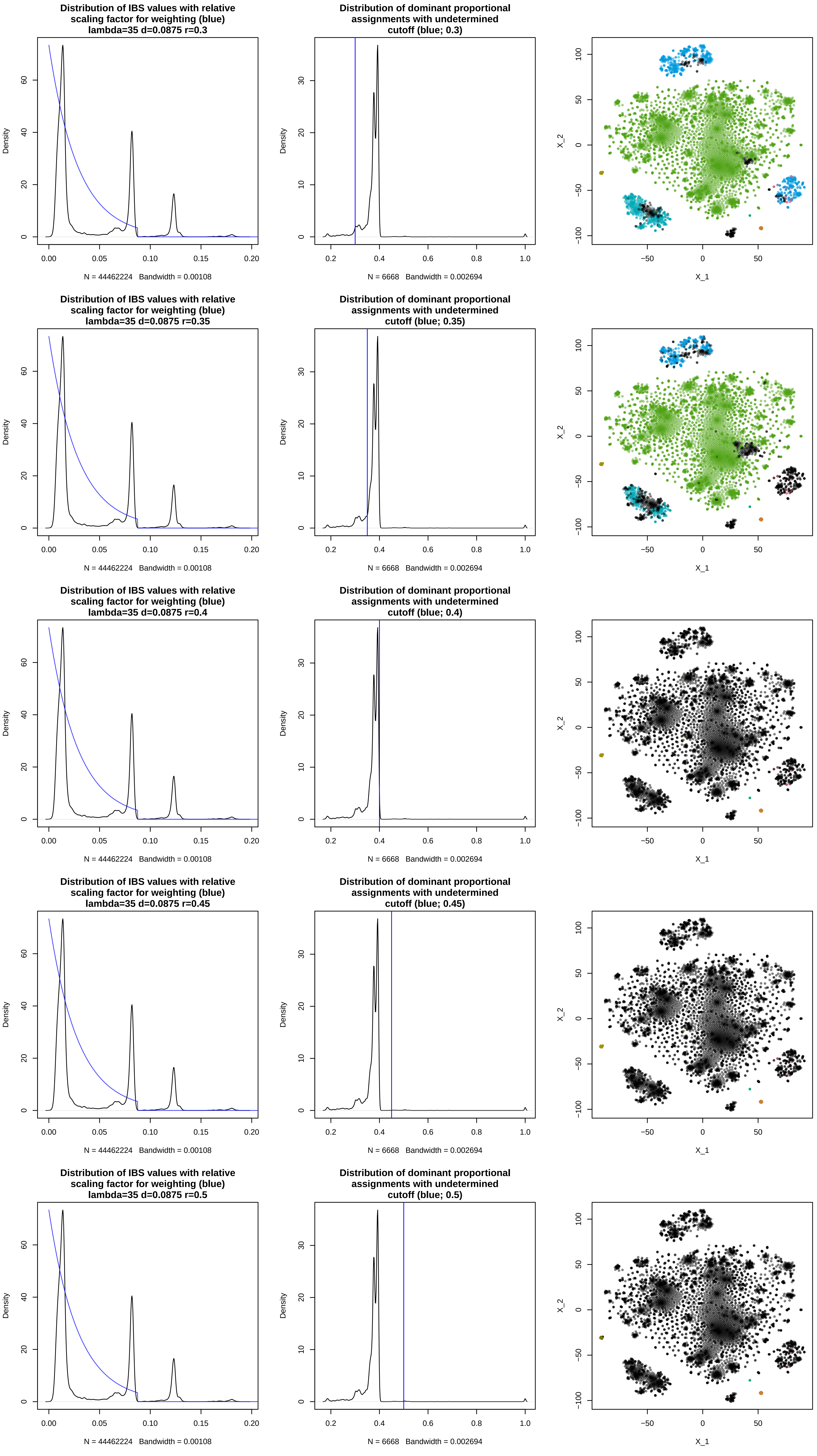

Distribution of IBS values with relative scaling factor for weighting (blue) lambda=35 d=0.0875 r=0.3
Density
N = 44462224 Bandwidth = 0.00108
Distribution of dominant proportional assignments with undetermined cutoff (blue; 0.3)
Density
N = 6668 Bandwidth = 0.002694
X_2
X_1
Distribution of IBS values with relative scaling factor for weighting (blue) lambda=35 d=0.0875 r=0.35
Density
N = 44462224 Bandwidth = 0.00108
Distribution of dominant proportional assignments with undetermined cutoff (blue; 0.35)
Density
N = 6668 Bandwidth = 0.002694
X_2
X_1
Distribution of IBS values with relative scaling factor for weighting (blue) lambda=35 d=0.0875 r=0.4
Density
N = 44462224 Bandwidth = 0.00108
Distribution of dominant proportional assignments with undetermined cutoff (blue; 0.4)
Density
N = 6668 Bandwidth = 0.002694
X_2
X_1
Distribution of IBS values with relative scaling factor for weighting (blue) lambda=35 d=0.0875 r=0.45
Density
N = 44462224 Bandwidth = 0.00108
Distribution of dominant proportional assignments with undetermined cutoff (blue; 0.45)
Density
N = 6668 Bandwidth = 0.002694
X_2
X_1
Distribution of IBS values with relative scaling factor for weighting (blue) lambda=35 d=0.0875 r=0.5
Density
N = 44462224 Bandwidth = 0.00108
Distribution of dominant proportional assignments with undetermined cutoff (blue; 0.5)
Density
N = 6668 Bandwidth = 0.002694
X_2
X_1

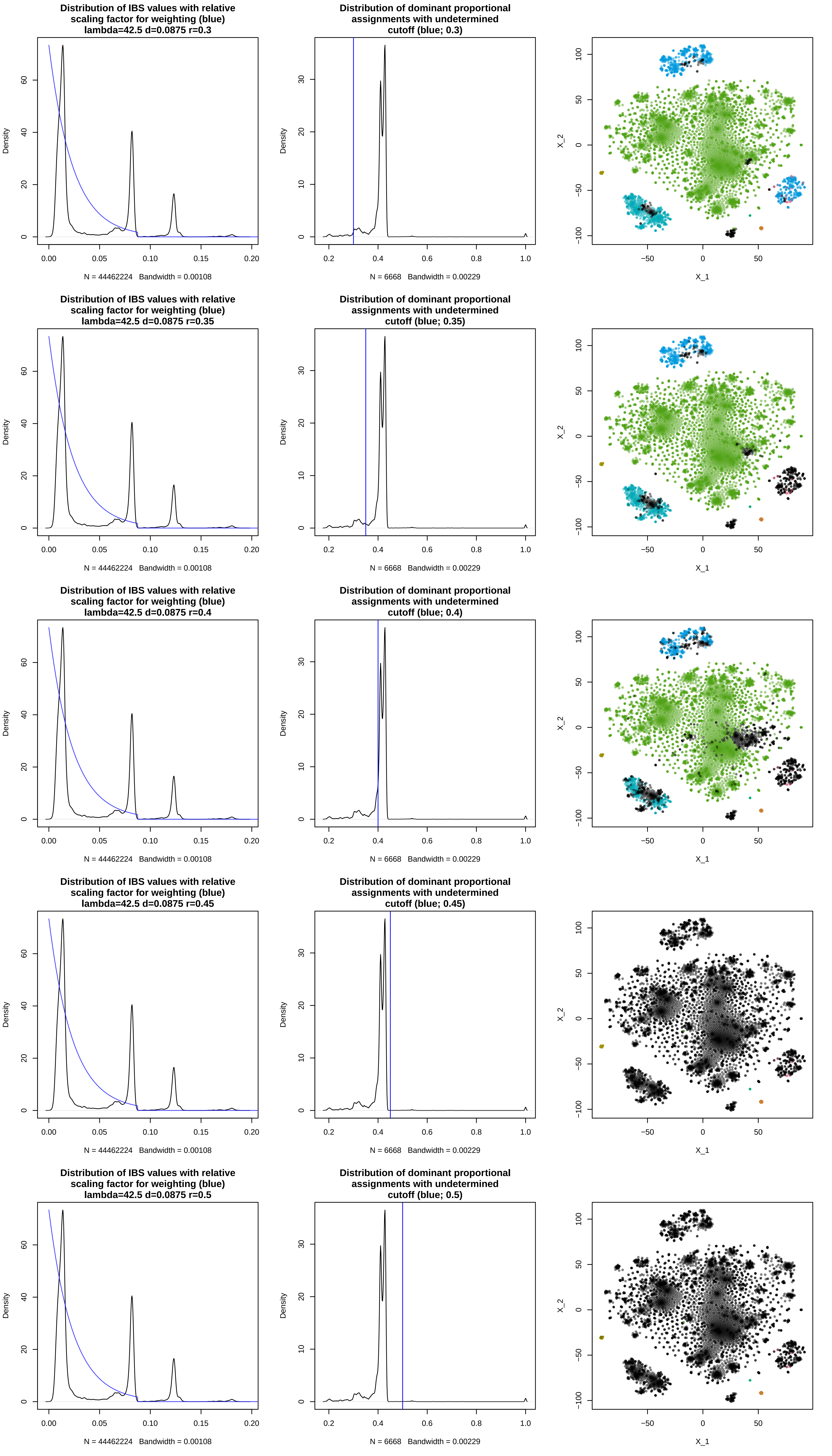
Distribution of IBS values with relative scaling factor for weighting (blue) lambda=42.5 d=0.0875 r=0.3
Density
N = 44462224 Bandwidth = 0.00108
Distribution of dominant proportional assignments with undetermined cutoff (blue; 0.3)
Density
N = 6668 Bandwidth = 0.00229
X_2
X_1
Distribution of IBS values with relative scaling factor for weighting (blue) lambda=42.5 d=0.0875 r=0.35
Density
N = 44462224 Bandwidth = 0.00108
Distribution of dominant proportional assignments with undetermined cutoff (blue; 0.35)
Density
N = 6668 Bandwidth = 0.00229
X_2
X_1
Distribution of IBS values with relative scaling factor for weighting (blue) lambda=42.5 d=0.0875 r=0.4
Density
N = 44462224 Bandwidth = 0.00108
Distribution of dominant proportional assignments with undetermined cutoff (blue; 0.4)
Density
N = 6668 Bandwidth = 0.00229
X_2
X_1
Distribution of IBS values with relative scaling factor for weighting (blue) lambda=42.5 d=0.0875 r=0.45
Density
N = 44462224 Bandwidth = 0.00108
Distribution of dominant proportional assignments with undetermined cutoff (blue; 0.45)
Density
N = 6668 Bandwidth = 0.00229
X_2
X_1
Distribution of IBS values with relative scaling factor for weighting (blue) lambda=42.5 d=0.0875 r=0.5
Density
N = 44462224 Bandwidth = 0.00108
Distribution of dominant proportional assignments with undetermined cutoff (blue; 0.5)
Density
N = 6668 Bandwidth = 0.00229
X_2
X_1

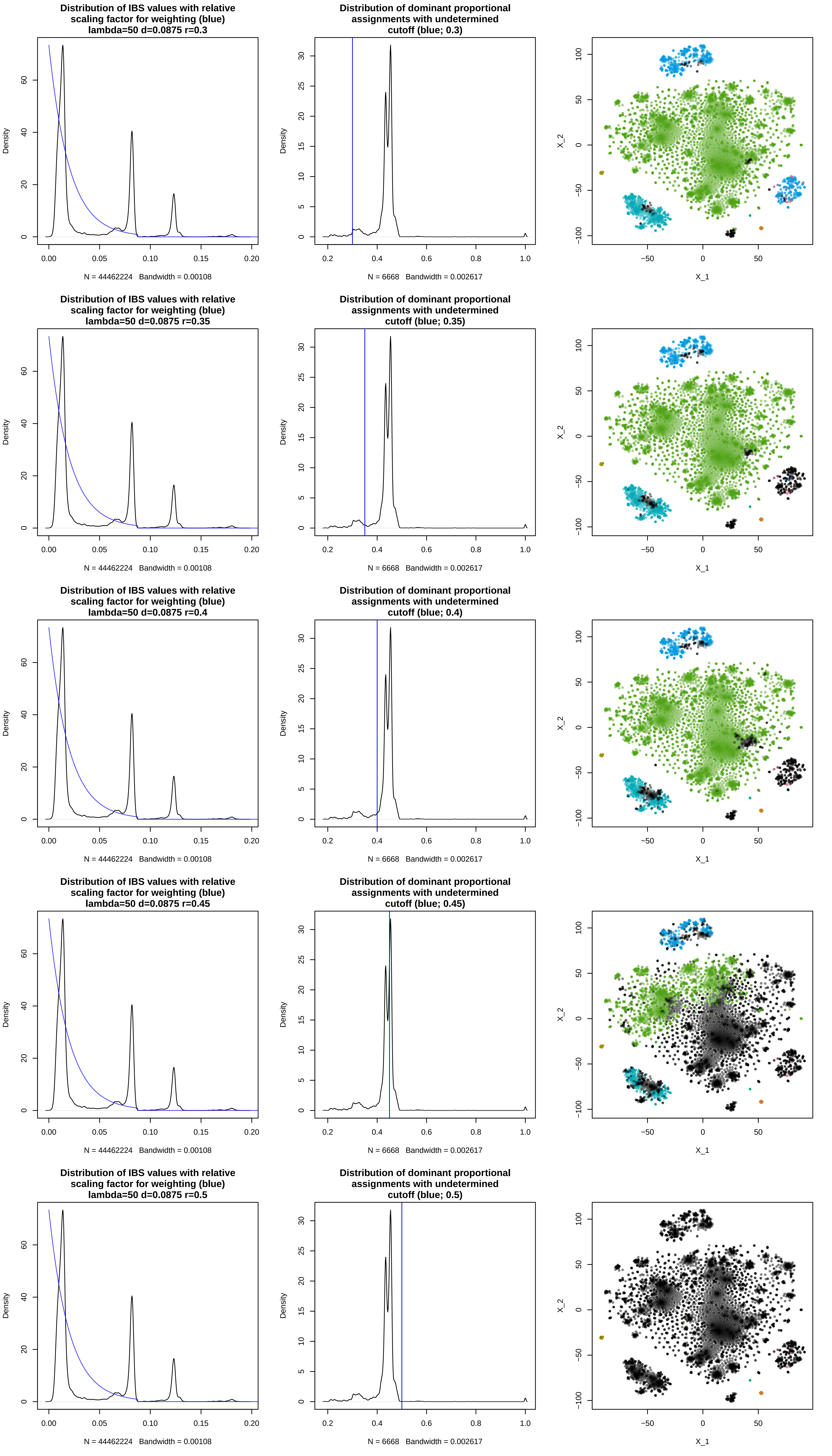
Distribution of IBS values with relative scaling factor for weighting (blue) lambda=50 d=0.0875 r=0.3
Distribution of dominant proportional assignments with undetermined cutoff (blue; 0.3)
Distribution of IBS values with relative scaling factor for weighting (blue) lambda=50 d=0.0875 r=0.35
Distribution of dominant proportional assignments with undetermined cutoff (blue; 0.35)
Distribution of IBS values with relative scaling factor for weighting (blue) lambda=50 d=0.0875 r=0.4
Distribution of dominant proportional assignments with undetermined cutoff (blue; 0.4)
Distribution of IBS values with relative scaling factor for weighting (blue) lambda=50 d=0.0875 r=0.45
Distribution of dominant proportional assignments with undetermined cutoff (blue; 0.45)
Distribution of IBS values with relative scaling factor for weighting (blue) lambda=50 d=0.0875 r=0.5
Distribution of dominant proportional assignments with undetermined cutoff (blue; 0.5)
Density
X_1
X_2
N = 44462224 Bandwidth = 0.00108
N = 6668 Bandwidth = 0.002617

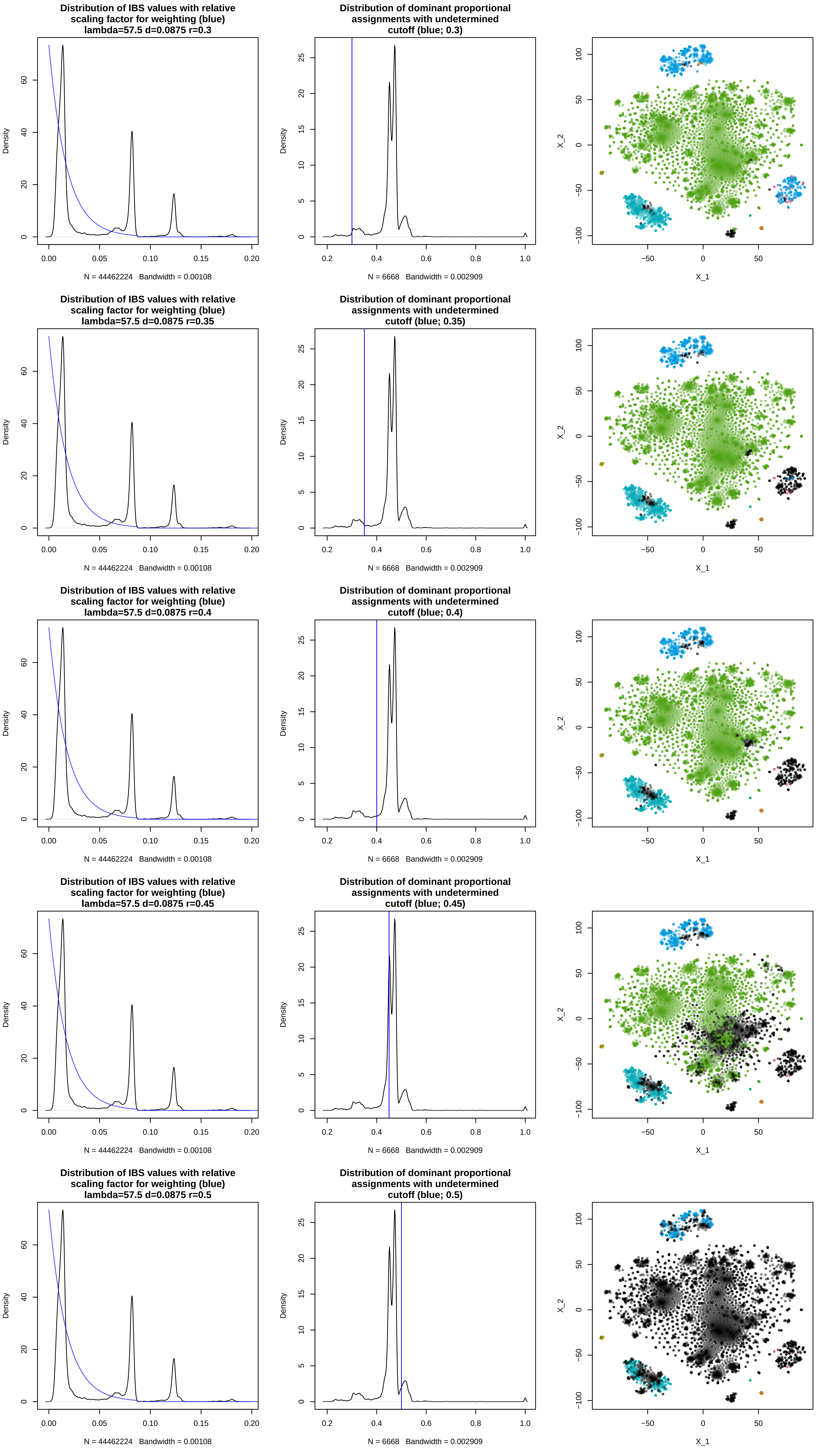
Distribution of IBS values with relative scaling factor for weighting (blue) lambda=57.5 d=0.0875 r=0.3
Distribution of dominant proportional assignments with undetermined cutoff (blue; 0.3)
Distribution of IBS values with relative scaling factor for weighting (blue) lambda=57.5 d=0.0875 r=0.35
Distribution of dominant proportional assignments with undetermined cutoff (blue; 0.35)
Distribution of IBS values with relative scaling factor for weighting (blue) lambda=57.5 d=0.0875 r=0.4
Distribution of dominant proportional assignments with undetermined cutoff (blue; 0.4)
Distribution of IBS values with relative scaling factor for weighting (blue) lambda=57.5 d=0.0875 r=0.45
Distribution of dominant proportional assignments with undetermined cutoff (blue; 0.45)
Distribution of IBS values with relative scaling factor for weighting (blue) lambda=57.5 d=0.0875 r=0.5
Distribution of dominant proportional assignments with undetermined cutoff (blue; 0.5)
Density
N = 44462224 Bandwidth = 0.00108
N = 6668 Bandwidth = 0.002909
X_1
X_2

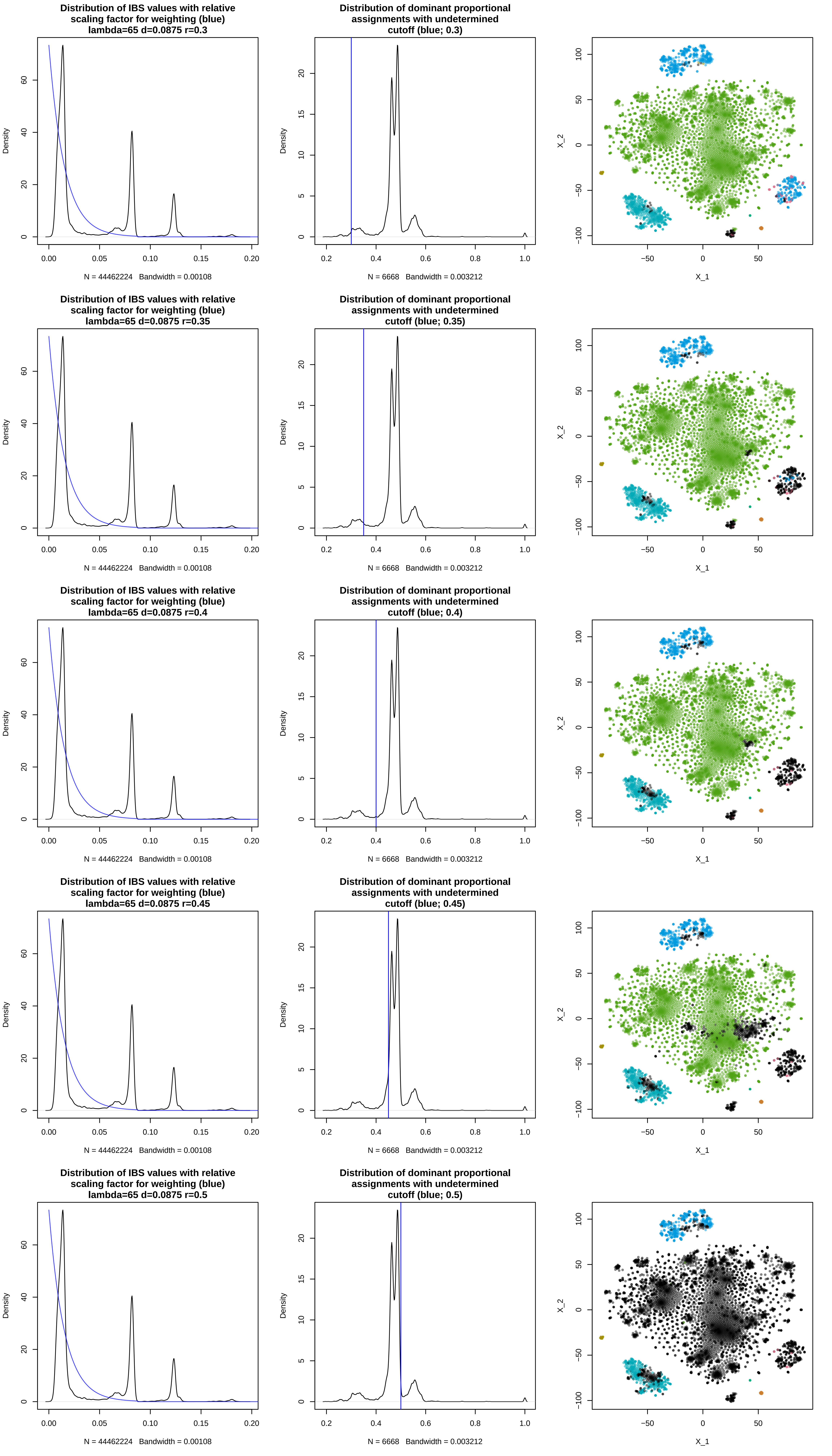

Distribution of IBS values with relative scaling factor for weighting (blue) lambda=65 d=0.0875 r=0.3
Density
N = 44462224 Bandwidth = 0.00108
Distribution of dominant proportional assignments with undetermined cutoff (blue; 0.3)
Density
N = 6668 Bandwidth = 0.003212
X_2
X_1
Distribution of IBS values with relative scaling factor for weighting (blue) lambda=65 d=0.0875 r=0.35
Density
N = 44462224 Bandwidth = 0.00108
Distribution of dominant proportional assignments with undetermined cutoff (blue; 0.35)
Density
N = 6668 Bandwidth = 0.003212
X_2
X_1
Distribution of IBS values with relative scaling factor for weighting (blue) lambda=65 d=0.0875 r=0.4
Density
N = 44462224 Bandwidth = 0.00108
Distribution of dominant proportional assignments with undetermined cutoff (blue; 0.4)
Density
N = 6668 Bandwidth = 0.003212
X_2
X_1
Distribution of IBS values with relative scaling factor for weighting (blue) lambda=65 d=0.0875 r=0.45
Density
N = 44462224 Bandwidth = 0.00108
Distribution of dominant proportional assignments with undetermined cutoff (blue; 0.45)
Density
N = 6668 Bandwidth = 0.003212
X_2
X_1
Distribution of IBS values with relative scaling factor for weighting (blue) lambda=65 d=0.0875 r=0.5
Density
N = 44462224 Bandwidth = 0.00108
Distribution of dominant proportional assignments with undetermined cutoff (blue; 0.5)
Density
N = 6668 Bandwidth = 0.003212
X_2
X_1

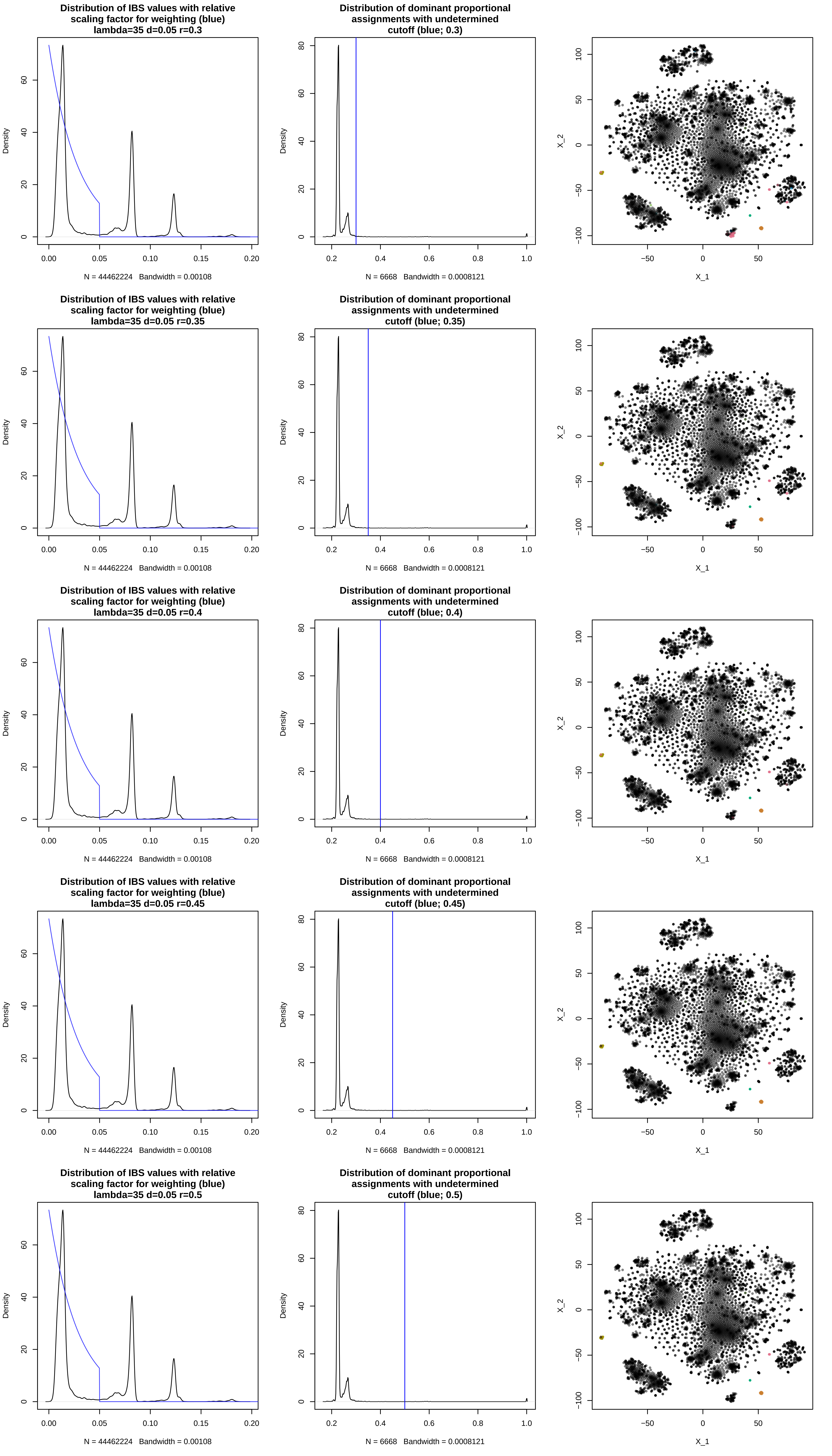
Distribution of IBS values with relative scaling factor for weighting (blue) lambda=35 d=0.05 r=0.3
Distribution of dominant proportional assignments with undetermined cutoff (blue; 0.3)
Distribution of IBS values with relative scaling factor for weighting (blue) lambda=35 d=0.05 r=0.35
Distribution of dominant proportional assignments with undetermined cutoff (blue; 0.35)
Distribution of IBS values with relative scaling factor for weighting (blue) lambda=35 d=0.05 r=0.4
Distribution of dominant proportional assignments with undetermined cutoff (blue; 0.4)
Distribution of IBS values with relative scaling factor for weighting (blue) lambda=35 d=0.05 r=0.45
Distribution of dominant proportional assignments with undetermined cutoff (blue; 0.45)
Distribution of IBS values with relative scaling factor for weighting (blue) lambda=35 d=0.05 r=0.5
Distribution of dominant proportional assignments with undetermined cutoff (blue; 0.5)
Density
N = 44462224 Bandwidth = 0.00108
N = 6668 Bandwidth = 0.0008121
X_1
X_2

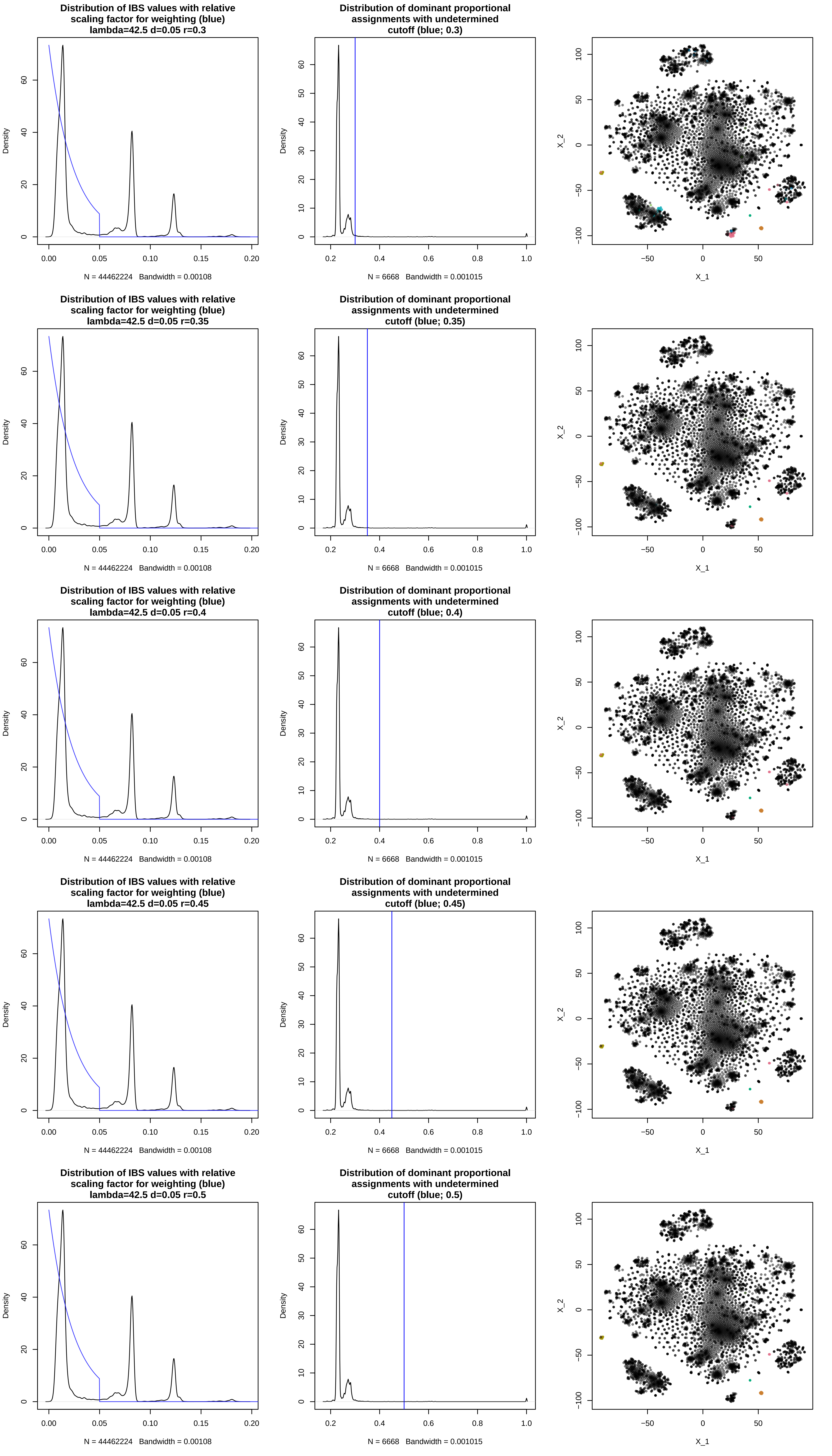
Distribution of IBS values with relative scaling factor for weighting (blue) lambda=42.5 d=0.05 r=0.3
Density
N = 44462224 Bandwidth = 0.00108
Distribution of dominant proportional assignments with undetermined cutoff (blue; 0.3)
Density
N = 6668 Bandwidth = 0.001015
X_1
X_2
Distribution of IBS values with relative scaling factor for weighting (blue) lambda=42.5 d=0.05 r=0.35
Density
N = 44462224 Bandwidth = 0.00108
Distribution of dominant proportional assignments with undetermined cutoff (blue; 0.35)
Density
N = 6668 Bandwidth = 0.001015
X_1
X_2
Distribution of IBS values with relative scaling factor for weighting (blue) lambda=42.5 d=0.05 r=0.4
Density
N = 44462224 Bandwidth = 0.00108
Distribution of dominant proportional assignments with undetermined cutoff (blue; 0.4)
Density
N = 6668 Bandwidth = 0.001015
X_1
X_2
Distribution of IBS values with relative scaling factor for weighting (blue) lambda=42.5 d=0.05 r=0.45
Density
N = 44462224 Bandwidth = 0.00108
Distribution of dominant proportional assignments with undetermined cutoff (blue; 0.45)
Density
N = 6668 Bandwidth = 0.001015
X_1
X_2
Distribution of IBS values with relative scaling factor for weighting (blue) lambda=42.5 d=0.05 r=0.5
Density
N = 44462224 Bandwidth = 0.00108
Distribution of dominant proportional assignments with undetermined cutoff (blue; 0.5)
Density
N = 6668 Bandwidth = 0.001015
X_1
X_2

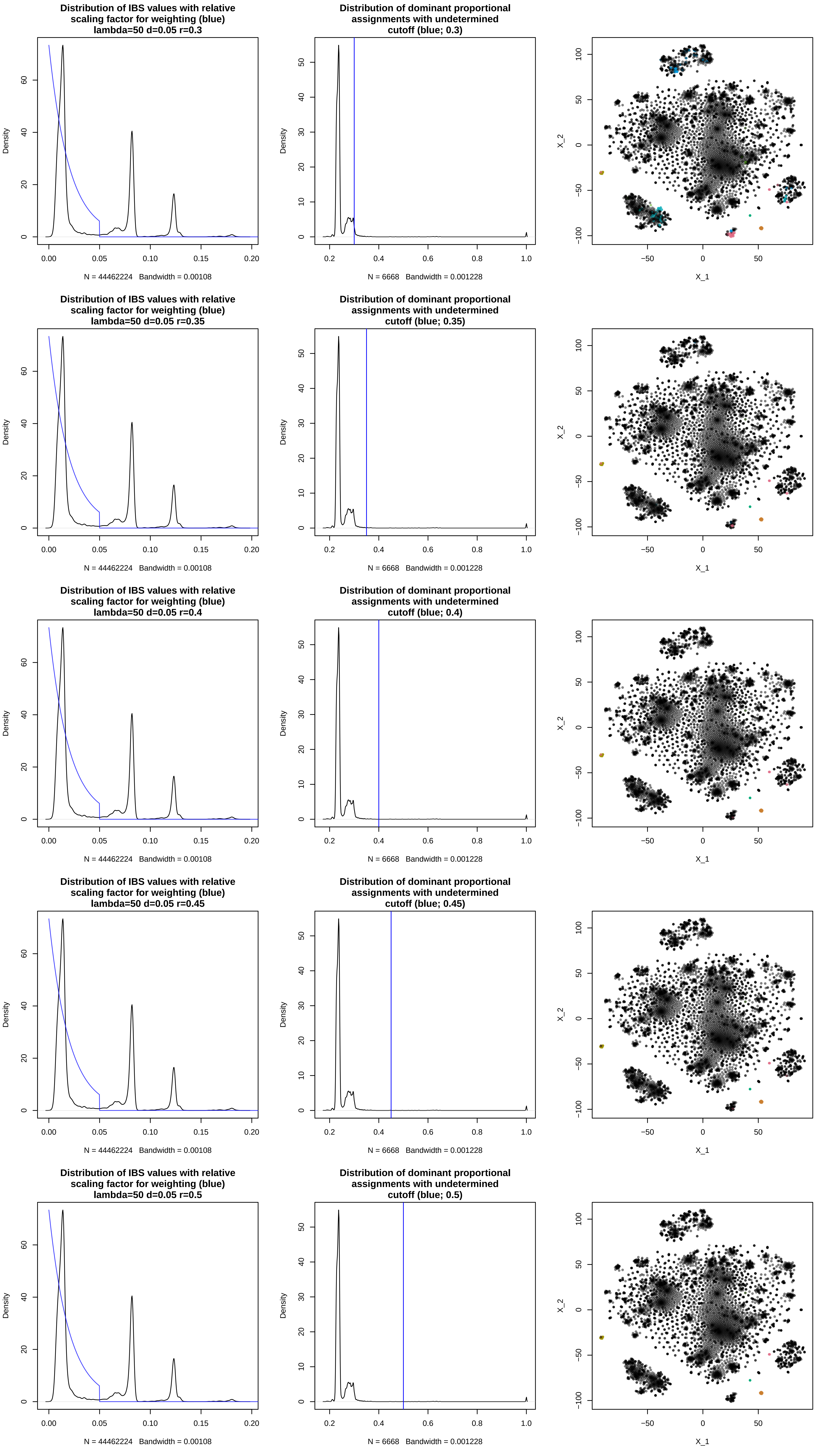
Distribution of IBS values with relative scaling factor for weighting (blue) lambda=50 d=0.05 r=0.3
Distribution of dominant proportional assignments with undetermined cutoff (blue; 0.3)
Distribution of IBS values with relative scaling factor for weighting (blue) lambda=50 d=0.05 r=0.35
Distribution of dominant proportional assignments with undetermined cutoff (blue; 0.35)
Distribution of IBS values with relative scaling factor for weighting (blue) lambda=50 d=0.05 r=0.4
Distribution of dominant proportional assignments with undetermined cutoff (blue; 0.4)
Distribution of IBS values with relative scaling factor for weighting (blue) lambda=50 d=0.05 r=0.45
Distribution of dominant proportional assignments with undetermined cutoff (blue; 0.45)
Distribution of IBS values with relative scaling factor for weighting (blue) lambda=50 d=0.05 r=0.5
Distribution of dominant proportional assignments with undetermined cutoff (blue; 0.5)
Density
N = 44462224 Bandwidth = 0.00108
N = 6668 Bandwidth = 0.001228
X_1
X_2

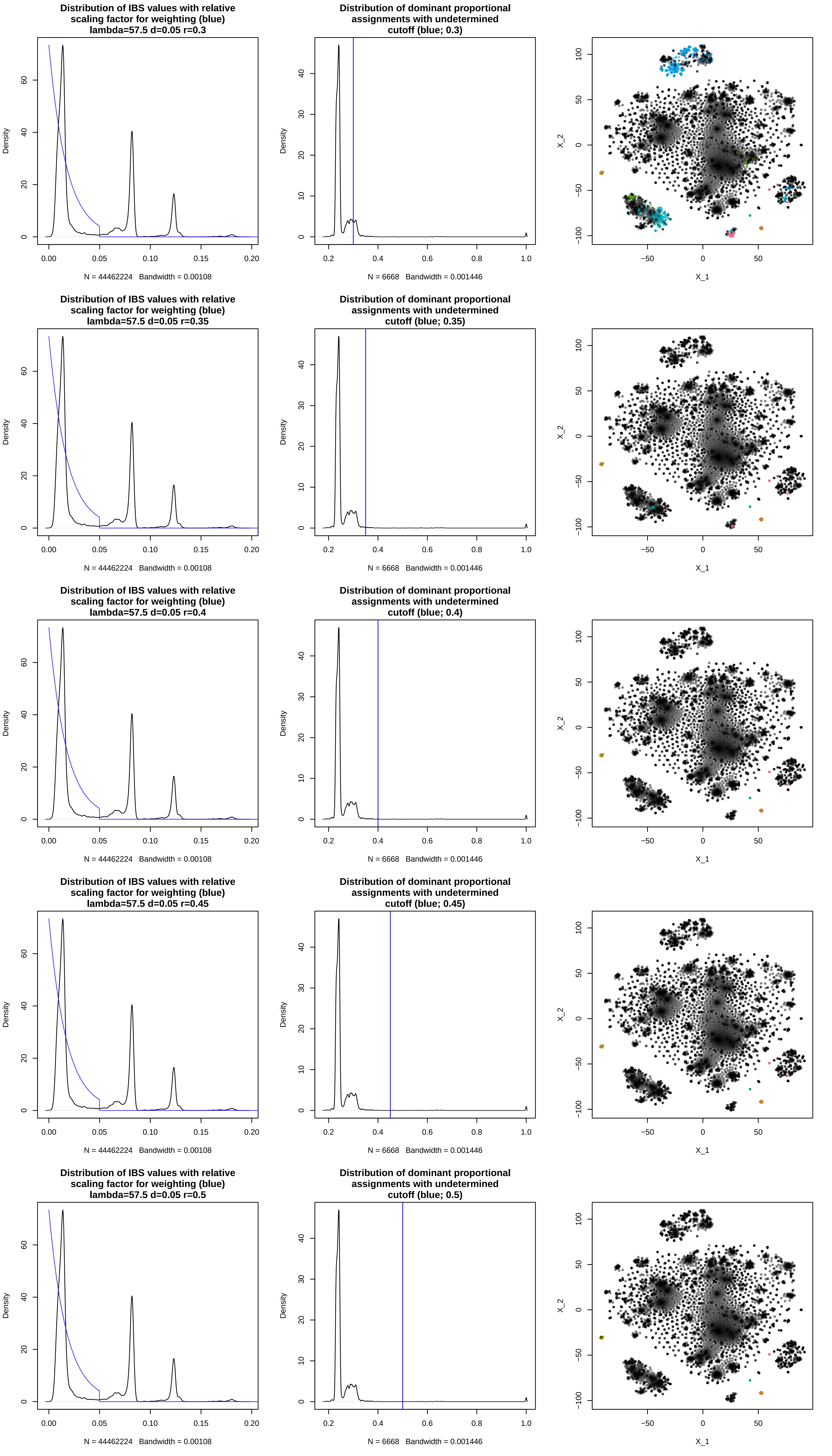
Distribution of IBS values with relative scaling factor for weighting (blue) lambda=57.5 d=0.05 r=0.3
Density
N = 44462224 Bandwidth = 0.00108
Distribution of dominant proportional assignments with undetermined cutoff (blue; 0.3)
Density
N = 6668 Bandwidth = 0.001446
X_1
X_2
Distribution of IBS values with relative scaling factor for weighting (blue) lambda=57.5 d=0.05 r=0.35
Density
N = 44462224 Bandwidth = 0.00108
Distribution of dominant proportional assignments with undetermined cutoff (blue; 0.35)
Density
N = 6668 Bandwidth = 0.001446
X_1
X_2
Distribution of IBS values with relative scaling factor for weighting (blue) lambda=57.5 d=0.05 r=0.4
Density
N = 44462224 Bandwidth = 0.00108
Distribution of dominant proportional assignments with undetermined cutoff (blue; 0.4)
Density
N = 6668 Bandwidth = 0.001446
X_1
X_2
Distribution of IBS values with relative scaling factor for weighting (blue) lambda=57.5 d=0.05 r=0.45
Density
N = 44462224 Bandwidth = 0.00108
Distribution of dominant proportional assignments with undetermined cutoff (blue; 0.45)
Density
N = 6668 Bandwidth = 0.001446
X_1
X_2
Distribution of IBS values with relative scaling factor for weighting (blue) lambda=57.5 d=0.05 r=0.5
Density
N = 44462224 Bandwidth = 0.00108
Distribution of dominant proportional assignments with undetermined cutoff (blue; 0.5)
Density
N = 6668 Bandwidth = 0.001446
X_1
X_2

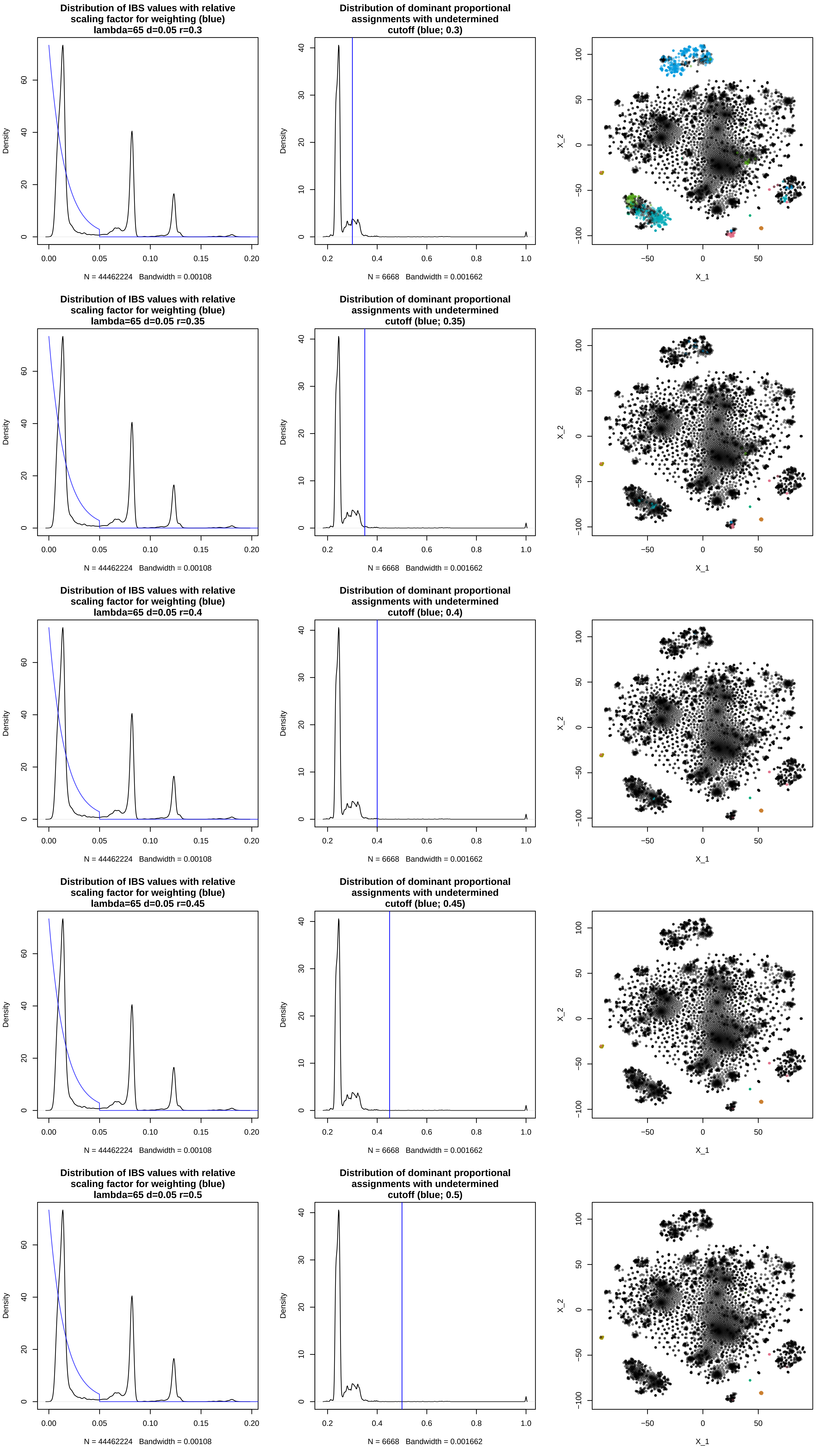

Distribution of IBS values with relative scaling factor for weighting (blue) lambda=65 d=0.05 r=0.3
Density
N = 44462224 Bandwidth = 0.00108
Distribution of dominant proportional assignments with undetermined cutoff (blue; 0.3)
Density
N = 6668 Bandwidth = 0.001662
X_2
X_1
Distribution of IBS values with relative scaling factor for weighting (blue) lambda=65 d=0.05 r=0.35
Density
N = 44462224 Bandwidth = 0.00108
Distribution of dominant proportional assignments with undetermined cutoff (blue; 0.35)
Density
N = 6668 Bandwidth = 0.001662
X_2
X_1
Distribution of IBS values with relative scaling factor for weighting (blue) lambda=65 d=0.05 r=0.4
Density
N = 44462224 Bandwidth = 0.00108
Distribution of dominant proportional assignments with undetermined cutoff (blue; 0.4)
Density
N = 6668 Bandwidth = 0.001662
X_2
X_1
Distribution of IBS values with relative scaling factor for weighting (blue) lambda=65 d=0.05 r=0.45
Density
N = 44462224 Bandwidth = 0.00108
Distribution of dominant proportional assignments with undetermined cutoff (blue; 0.45)
Density
N = 6668 Bandwidth = 0.001662
X_2
X_1
Distribution of IBS values with relative scaling factor for weighting (blue) lambda=65 d=0.05 r=0.5
Density
N = 44462224 Bandwidth = 0.00108
Distribution of dominant proportional assignments with undetermined cutoff (blue; 0.5)
Density
N = 6668 Bandwidth = 0.001662
X_2
X_1
