## Supplementary figures and images for "Genebank genomics allows greatly improved taxonomic correction for *Capsicum spp*. accessions using a novel automated classification method"

### Supplemental figures 2

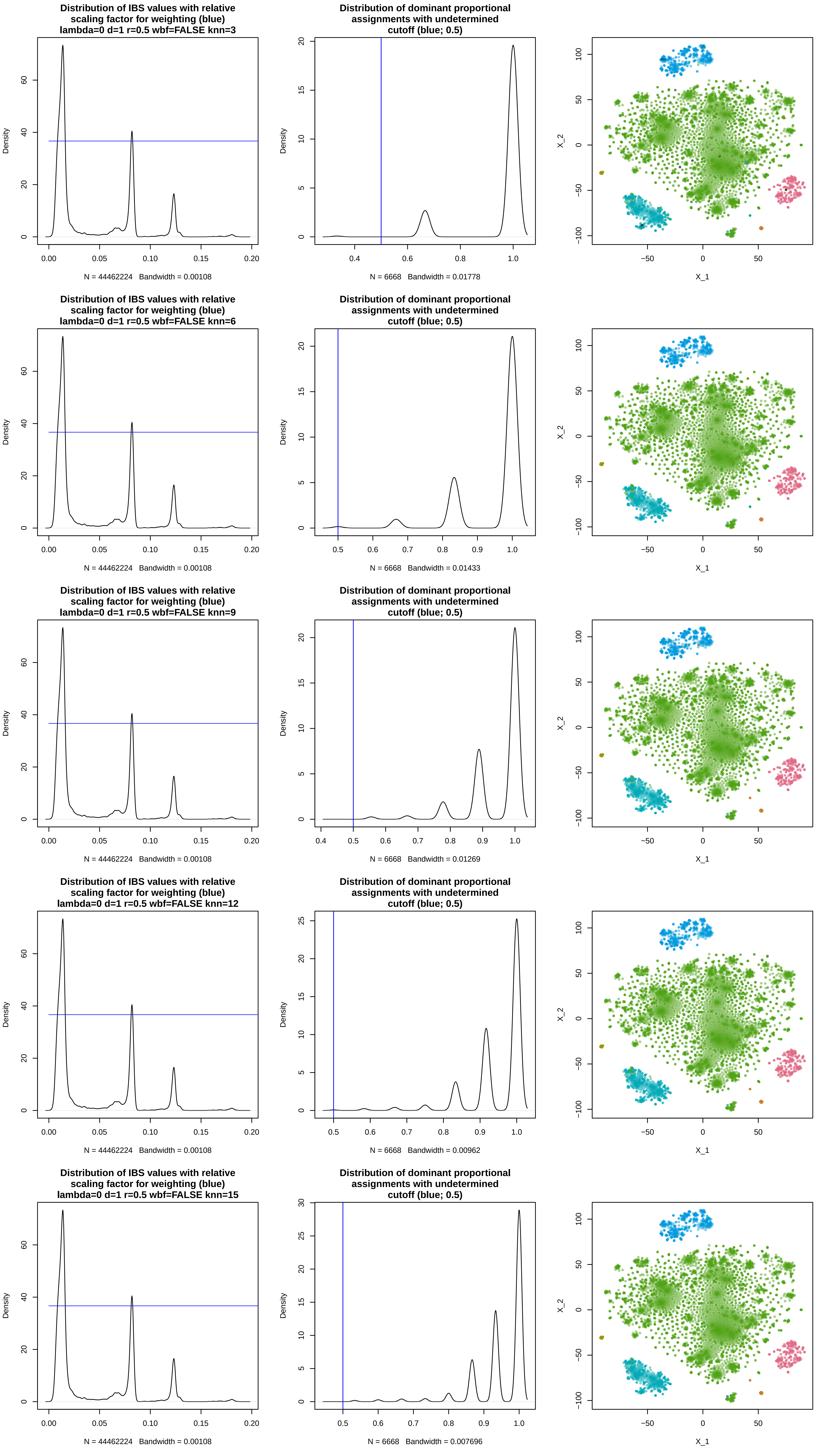

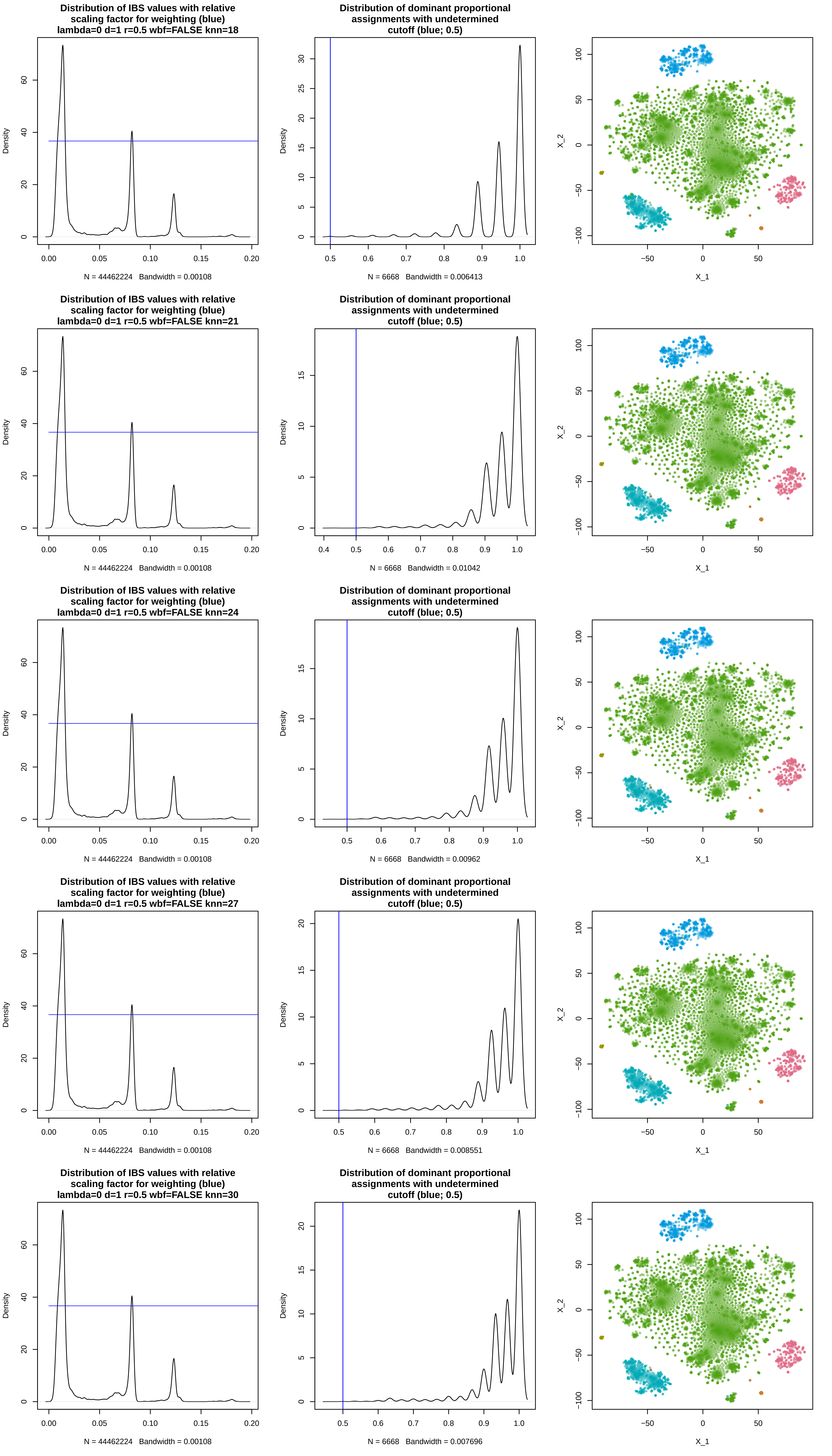

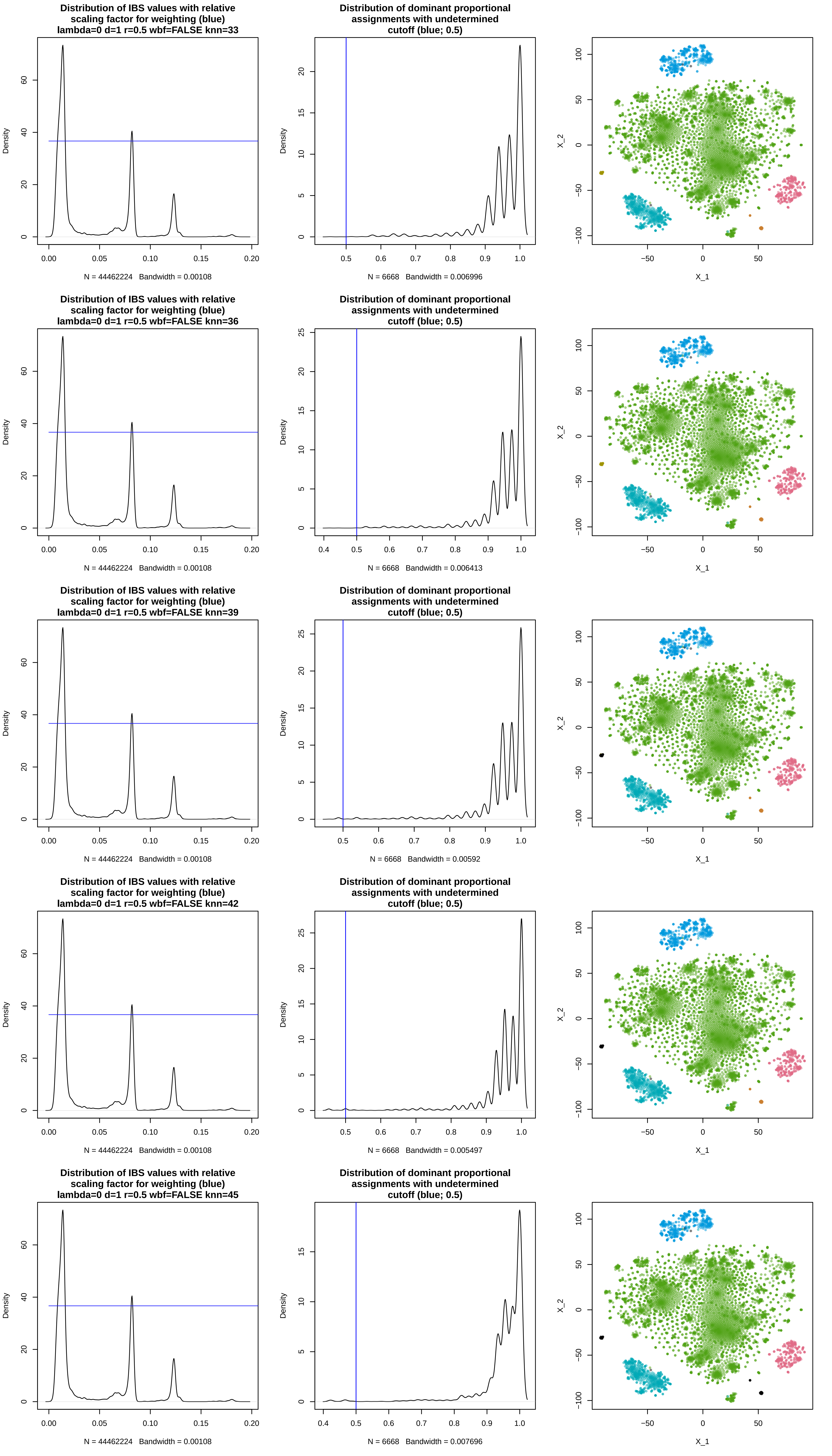

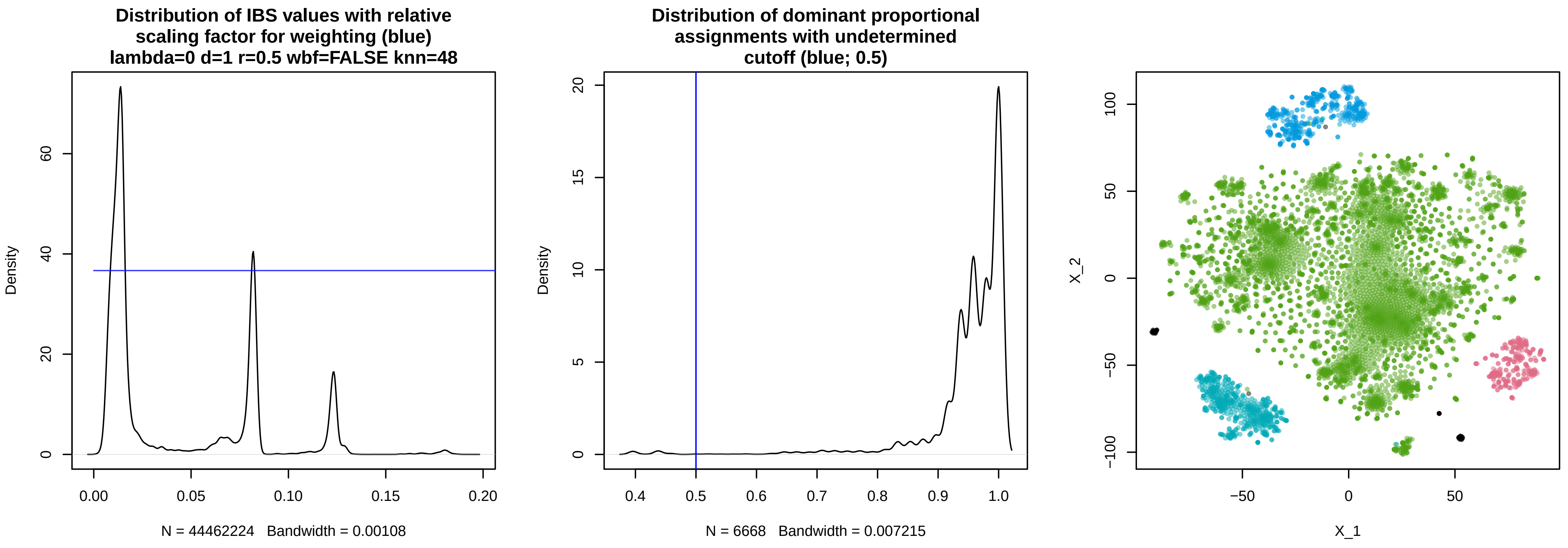
